# Midbrain *Tet1* dosage defines inter-individual binge-eating susceptibility

**DOI:** 10.64898/2026.03.14.711800

**Authors:** Tim Gruber, Robert A. Chesters, Luca Fagnocchi, Xinyang Yu, Zhen Fu, Kristin Gallik, Heiko Backes, Robert Vaughan, Melanie Huber, Meri De Angelis, Josef Gullmets, Holly Dykstra, Stefanos Apostle, Taylor Cook, Justin Kulchycki, Lisa DeCamp, Timo D. Müller, Katharina Timper, the PERMUTE,IMAGEN and ESTRA consortia, Sylvane Desrivières, Rachel N. Lippert, J. Andrew Pospisilik

**Affiliations:** Department of Epigenetics, Van Andel Institute, Grand Rapids, MI, USA; Department of Metabolism and Nutritional Programming, Van Andel Institute, Grand Rapids, MI, USA; German Center for Diabetes Research (DZD), Neuherberg, Germany; Neurocircuit Development and Function, German Institute for Nutrition (DIfE), Potsdam-Rehbrücke, Germany; Social, Genetic and Developmental Psychiatry Centre, Institute of Psychiatry, Psychology and Neuroscience, King’s College London, London, UK; Bioinformatics and Biostatistics Core Facility, Van Andel Institute, Grand Rapids, MI, USA; Optical Imaging Core Facility, Van Andel Institute, Grand Rapids, MI, USA; Max Planck Institute for Metabolism Research, Cologne, Germany; Institute for Diabetes and Obesity, Helmholtz Diabetes Center at Helmholtz Munich, Neuherberg, Germany; Institute of Experimental Genetics, Helmholtz Center Munich, German Research Center for Environmental Health, Neuherberg, Germany; Walther-Straub-Institute of Pharmacology and Toxicology, Ludwig-Maximilians-Universität München (LMU Munich), Germany; Institute for Translational Metabolism Research, Helmholtz Diabetes Center at Helmholtz Munich, Neuherberg, Germany; Institute for Clinical Nutritional Medicine, Technical University of Munich, TUM School of Medicine and Health, TUM University Hospital, TUM Campus at the Olympiapark, Munich, Germany; Else Kröner-Fresenius-Center for Nutritional Medicine, Technical University of Munich, Freising, Germany; Department Biomedicine Basel, University Hospital of Basel and University of Basel, Basel, Switzerland

## Abstract

Binge-eating disorder (BED) is the most common eating disorder worldwide and carries life-altering comorbidities. While genetic and environmental risk factors have been identified, the mechanisms that determine inter-individual susceptibility to BED remain largely unknown. Here, we demonstrate that developmental dosage of the DNA hydroxymethylase *Tet1* defines stable inter-individual differences in binge-eating susceptibility. In mice, midbrain dopaminergic neurons of the ventral tegmental area (VTA^DA^) are essential for the induction of addictive binge-eating behavior, express high levels of *Tet1*, and undergo rapid and widespread DNA hydroxymethylation remodeling upon experimental binge eating. Strikingly, *Tet1* haploinsufficiency creates pronounced inter-individual variation in binge-eating susceptibility even among genetically identical mice, which we trace to reduced connectivity between the prelimbic medial prefrontal cortex (mPFC^PL^) and the VTA. Chemogenetic inhibition of mPFC^PL^→VTA projections reduces binge-eating susceptibility, whereas EGR1-guided re-activation of TET1 in VTA dopaminergic neurons restores susceptibility, supporting a causal role for this axis. Importantly, *TET1* promoter methylation in patients associates with binge-eating behavior and reward-circuit function, suggesting conservation of this regulatory network in humans. Collectively, these findings identify *Tet1* dosage as a novel regulator of binge-eating susceptibility and provide a mechanistic basis for how inter-individual differences in behavior are established.

## Introduction

Binge-eating disorder (BED) is the most common eating disorder worldwide, affecting 2–3 % of the general population.^1,2^ BED was officially recognized as a mental disorder in 2013 (DSM-5) and is characterized by compulsive and rapid consumption of large amounts of energy-dense food (i.e., recurrent ‘binge episodes’). The disorder is associated with serious metabolic and psychiatric comorbidities including obesity, diabetes, depression, anxiety, and low self-esteem.^3–5^ BED shares clinical, genetic, and neurobiological features with impulse control and addiction-related disorders, including substance use disorder, pathological gambling, and compulsive buying.^6–8^ BED, however, is unique because its primary reward stimulus, food, is essential for survival.

Mammals have evolved numerous neurocircuits to control eating behavior. Included, midbrain dopamine neurons^9–11^ in the ventral tegmental area (VTA^DA^ neurons) reinforce eating (and other behaviors) by encoding reward-related predictions and learning.^12^ VTA^DA^ neurons project to key limbic and cortical targets, including the nucleus accumbens (NAc) and medial prefrontal cortex (mPFC), where cue-dependent dopamine release promotes reward-contingent learning, the perceived value of cues, and motivated behaviors.^13^ Unsurprisingly, these same pathways are implicated in a variety of addictions.^14,15^ Feedback from higher-order regions such as the striatum and mPFC finetune VTA^DA^ excitability.^16–21^

Despite the ubiquity of high-reward, calorie-dense foods in the developed world, most people resist binge eating. That said, up to 40 % of overweight individuals do experience food cravings to the point of losing control over food intake.^22^ Twin and adoption studies estimate the heritability of BED at 39–45 % indicating a substantial role for both genetic and non-genetic influences.^23–25^ To date, no diagnostic effectively identifies or predicts individuals at risk. Interestingly, pronounced differences in addiction- and reward-related behaviors are observed between monozygotic twins^26–28^ and between group-housed inbred (genetically ‘identical’) rodents.^29–31^ These findings suggest that stochastic or epigenetic processes might calibrate inter-individual reward setpoints^32,33^ defined as intrinsic triggering-levels for motivational states. Indeed, a central challenge in behavioral neuroscience is to identify the molecular determinants of interindividual differences in behavioral setpoints, i.e., the origins of individuality itself.^34,35^

Here, we identify the DNA hydroxymethylase TET1^36,37^ as a critical regulator of binge-eating susceptibility. We show that *Tet1* shapes inter-individual vulnerability by influencing both the developmental establishment of binge behavior setpoints, and locus-specific epigenome regulation in adult VTA^DA^ neurons. This *Tet1*-dependent regulatory circuitry, mapped using mouse transgenic systems, appears conserved. In humans, *TET1* promoter methylation associates with reward-circuit function and binge eating in patients and shows evidence of genotype dependence. Collectively, these findings identify *Tet1* dosage as an important regulator of VTA^DA^ circuit development and function, and one of the first known molecular determinants of eating-behavior individuality.

## Results

### VTA^DA^ activity is necessary for eBED

VTA^DA^ neurons are critical regulators of reward processing and behavioral reinforcement,^11,12,30^ yet their involvement in BED had not been directly tested. Towards these ends, we generated a Ca^2+^-reporter mouse line (DAT-*ires*-Cre; AAV-FLEX-GCaMP6s) that enabled real-time monitoring of dopaminergic neuron activity via fiber photometry (**Figure 1A, 1B** and **Supplementary Figure 1A**). We stereotaxically injected AAV-FLEX-GCaMP6s into the VTA of ∈8 w old mice, implanted fiber-optic cannulae, allowed three weeks for recovery, and acclimated the animals for five days to the recording set-up. To model BED-like eating,^38,39^ we used an intermittent exposure paradigm, referred to here as ‘eBED’ (experimental BED). Mice are provided ad libitum access to chow diet in addition to a 2-hour daily binge opportunity, comprising access to high-fat diet (HFD; 60 % kcal). Intermittent HFD access is provided for five consecutive days during the peak of the circadian eating cycle. Controls (cHFD) instead receive continuous access to both HFD and chow (**Figure 1C**). Consistent with the goals of the model, eBED challenged animals progressively increase their HFD intake during the binge opportunity, to the point that they consumed almost half of their daily calories within the short 2 h window (**Figure 1D**). This escalation of HFD intake was driven by voracious eating during the experimental binge, with observed increases in both eating rate (**Figure 1E**) and meal size (**Figure 1F**). By contrast, cHFD controls showed no escalation, required more than three times as long to reach comparable HFD intake levels (6.4 ± 0.7 *h*), and consumed HFD steadily throughout the day. Thus, we established a system for probing VTA^DA^ activity during binge-like eating.

**Figure 1.**
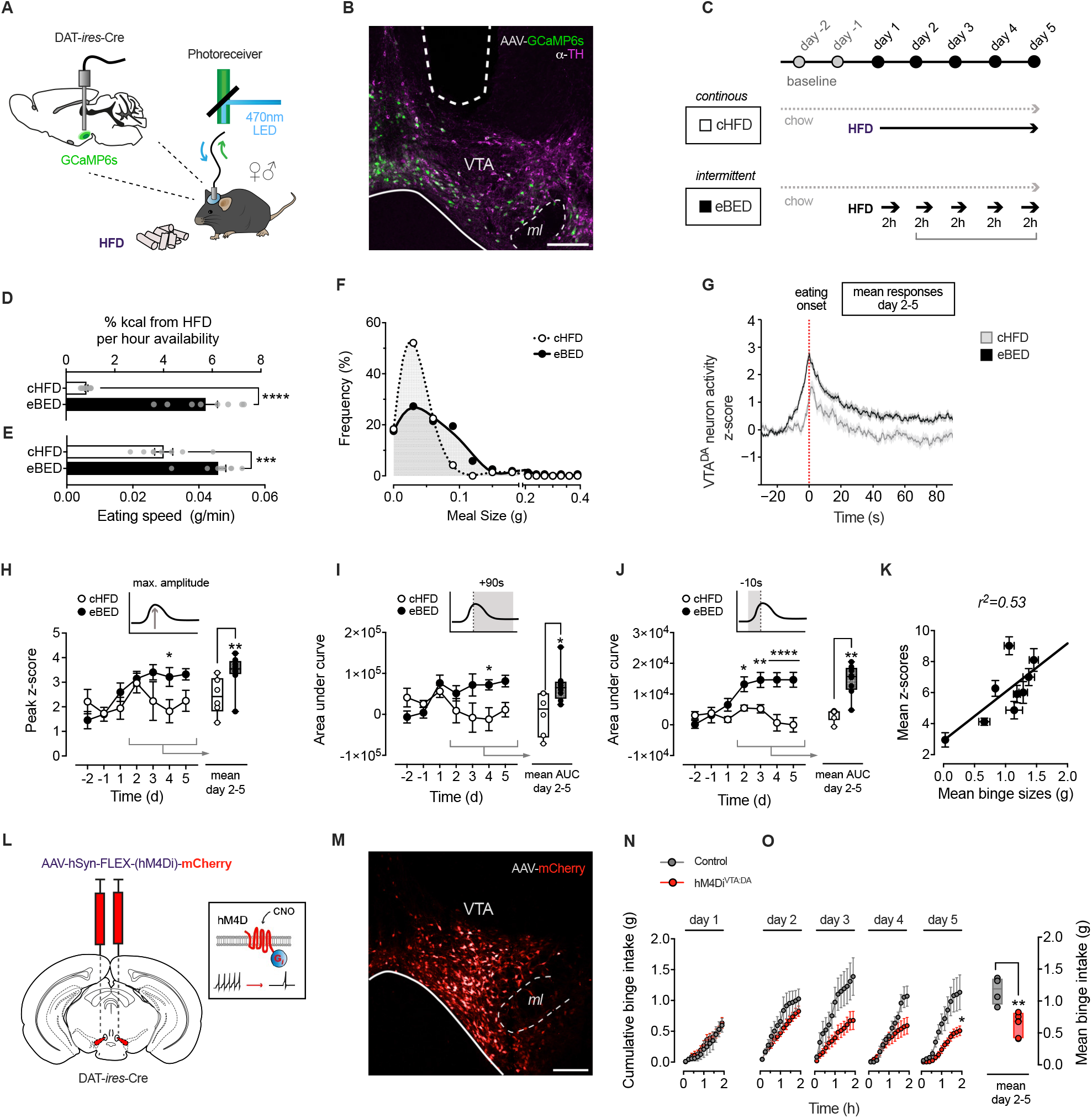
Escalatory VTA^DA^ activity is triggered by eBED and required for binge eating in mice. **(A)** Fiber photometry set up to assess real-time VTA^DA^ neuron activity in behaving mice. **(B)** GCaMP6s expression (green) colocalizing with VTA^DA^ neurons (TH^+^; magenta) and optic fiber tract. **(C)** Experimental overeating paradigm comparing five days of eBED (intermittent, 2 h-limited HFD access) to controls (continuous or cHFD); ad libitum chow diet was provided throughout. **(D)** Percent daily kilocalories from HFD per hours of availability. Data are presented as mean ± SEM. **** *P <* 0.0001. *n* = 8 mice (unpaired Student’s *t* -test). (E) Eating speed during HFD intake. Data are presented as mean ± SEM. *** *P <* 0.001. *n* = 8 mice (unpaired Student’s *t* -test). (F)Meal size distribution during 2h HFD access. *n* = 8 mice. **(G)** Fiber photometry VTA^DA^ recordings of mean z-score (day 2–5) at eating onset (0 s). Data are presented as mean ± SEM. *n* = 6–8 mice. **(H)** Peak z-score (maximum amplitude) per individual days (left panel) and mean values (day 2–5; right panel). Data are presented as mean ± SEM mean and minimum to maximum values. * *P <* 0.05, ** *P <* 0.01. *n* = 6–8 mice (multiple unpaired Student’s *t* -test). **(I)** Post-eating AUC (0–90s) of z-scores per individual days (left panel) and mean values (day 2–5; right panel). Data are presented as mean ± SEM and mean minimum to maximum values. * *P <* 0.05, ** *P <* 0.01. *n* = 6–8 mice (multiple unpaired Student’s *t* -test). **(J)** Pre-eating AUC (-10–0s) of z-scores per individual days (left panel) and mean values (day 2–5; right panel). Data are presented as mean ± SEM and mean minimum to maximum values. * *P <* 0.05, ** *P <* 0.01, **** *P <* 0.001. *n* = 6–8 mice (multiple unpaired Student’s *t* -test). **(K)** Linear regression of mean binge sizes and z-scores (10 min after HFD access; day 2–5). **(L)** Schematic illustrating VTA^DA^ neuron inhibition by virally expressed chemogenetic actuator hM4Di following injection of CNO (clozapine-N-oxide). **(M)** Confocal micrograph target validation of virally expressed mCherry in the VTA. **(N)** Cumulative binge intake of hM4Di^VTA:DA^ mice and controls on the first day of exposure. Data are presented as mean ± SEM. *n* = 4–7 mice (multiple unpaired Student’s *t* -test). **(O)** Cumulative binge intake of hM4Di^VTA:DA^ mice and controls per individual days (left panel) and mean values (day 2–5; right panel). Data are presented as mean ± SEM and minimum to maximum values. * *P <* 0.05, ** *P <* 0.01. *n* = 4–7 mice (multiple unpaired Student’s *t* -test).

Using this system, we characterized neuronal dynamics at eBED onset. Whereas interactions with chow triggered only modest VTA^DA^ responses (Days −2 and −1; **Supplementary Figure 1B**), we found that animals interacting for their first time with HFD pellets (whether in the eBED or control cHFD group) displayed robust VTA^DA^ activation, characterized by high and exceptionally long GCaMP6s responses (Day +1; **Supplementary Figure 1C-D**). These findings are consistent with a positive novelty response (also known as a reward-prediction error). Interestingly, we observed a pronounced divergence in VTA^DA^ responses between groups, from the second day onwards. Importantly, on the subsequent Days 2–5, VTA^DA^ activity during HFD-eating diverged between groups; cHFD controls showed transient, modest signals, whereas eBED animals exhibited persistently exaggerated responses (**Figure 1G**). These subsequent VTA^DA^ responses were shorter in duration than Day 1, but significantly stronger in maximum amplitude relative to cHFD controls (**Figure 1H**). Cumulative VTA^DA^ activity during eating was significantly higher in eBED compared to cHFD animals (AUC 0–90s; **Figure 1I**). Particularly striking, eBED mice exhibited substantial VTA^DA^ activation *preceding* their interaction with the HFD-pellet (**Figure 1J**; Days 2–5). This suggested that eBED induces ‘craving’-like neurobehavior, a common feature of impulse control disorders. Lastly, individual binge sizes were strongly correlated with GCaMP6s responses across eBED mice, indicating exceptionally tight coupling between VTA^DA^ activity and eating behavior. Thus, eBED triggers robust, escalatory VTA^DA^ neuron activation.

Finally, we tested whether these VTA^DA^ dynamics were required for the induction of hallmark binge-eating behaviors. We targeted the chemogenetic actuator, hM4Di, to VTA^DA^ neurons to enable their selective, on-demand inhibition, and compared binge eating in VTA^DA^-inhibited (hM4Di^VTA:DA^) versus control virus-injected mice. Specifically, DAT-*ires*-Cre mice were injected bilaterally with AAV-FLEX-hM4Di-mCherry virus (or control AAV-FLEX-mCherry)(**Figure 1L** and **1M**). Animals were allowed to recover for three weeks, acclimated to the experimental conditions, and then subject to the same eBED protocol as above. Both groups received the hM4Di-activating (VTA^DA^ inhibiting) designer ligand CNO (clozapine-N-oxide) 15 min before binge exposure. Interestingly, HFD intake on the first exposure day was equal between VTA^DA^ -inhibited animals and controls (**Figure 1N**), suggesting that novelty-associated eating is not sensitive to VTA^DA^ activation. Importantly however, while control mice showed the expected escalatory binge response over five days, hM4Di^VTA:DA^ mice were protected, taking in only modest amounts of HFD and showing no escalation in 2h HFD intake (**Figure 1O**). Consistent with these observations, hM4Di^VTA:DA^ animals showed no evidence of increased binge meal size or eating rate (**Supplementary Figure 1F** and **1G**). Noteworthy, daily caloric intake and locomotor activity were the same between groups, arguing against potentially confounding hypoactivity, bradykinesia, or differential energy balance (**Supplementary Figure 1H** and **1I**). Thus, VTA^DA^ neuronal activity is necessary for the induction of eBED.

### eBED triggers rapid rewiring of the VTA^DA^ epigenome

There remains an ongoing debate as to whether BED is a *bona fide* addiction.^40–43^ BED’s abrupt onset, its *binge-withdrawal-craving* cycles, and high recidivism are shared with other addictions and suggest an underlying molecular triggering process.^2^ Mounting evidence suggests that epigenetic programs can support such rapid remodeling of reward learning and compulsivity.^33,44^ We therefore asked whether eBED triggers addiction-like binge behavior through epigenetic rewiring of VTA^DA^ neurons. We profiled DNA methylation (5mC) and DNA *hydroxy*methylation (5hmC)^45–47^ in purified VTA^DA^ nuclei from mice following a modified eBED paradigm (eliminating HFD novelty with 4-weeks HFD pre-exposure)(**Figure 2A** and **Supplementary Figure 2A**).

**Figure 2.**
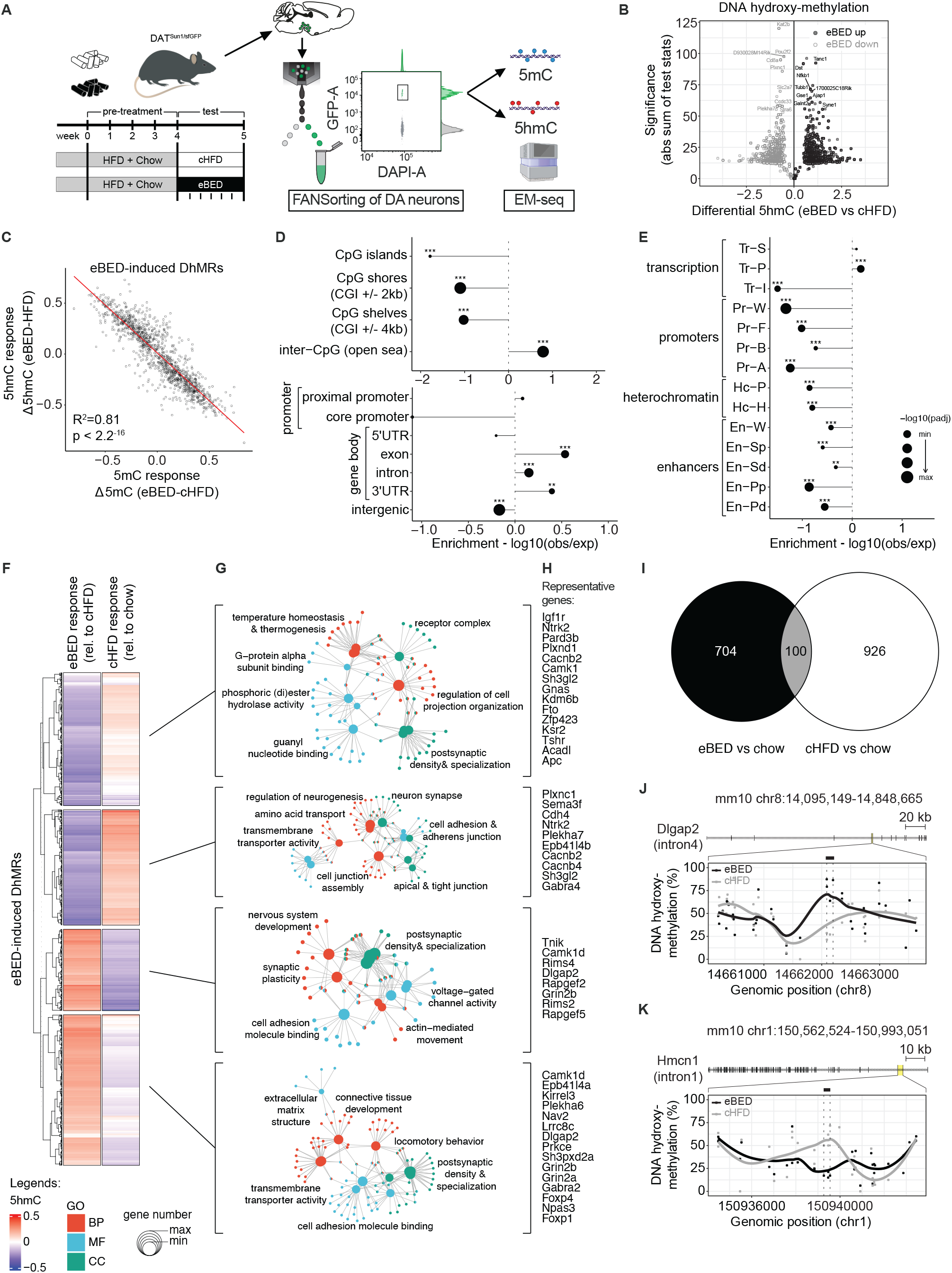
eBED triggers rapid and profound rewiring of the VTA^DA^ neuron epigenome. **(A)** Experimental paradigm of DAT^Sun1/sfGFP^ reporter mice with additional pre-exposure to HFD, FANSorting protocol (including representative gating plot) followed by Enzymatic Methyl (EM)-seq. *Created in BioRender.com*. **(B)** Volcano plot of DhMRs comparing eBED versus cHFD. **(C)** Scatter plot of 5hmC versus 5mC responses to eBED, relative to cHFD, on eBED-induced DhMRs. *R*^2^ and *p*-value from Pearson’s correlation. **(D)** Dotplot showing the relative enrichments and depletions of indicated genomic regions over eBED-induced DhMRs. *P*-values from Fisher’s exact test. **(E)** Dotplot showing the relative enrichments and depletions of indicated genomic regions identified at P0 age in mouse midbrain cells, over eBED-induced DhMRs. *P*-values from Fisher’s exact test. **(F)** Heatmap showing z-scored 5hmC signal on k-means clustering of eBED-induced DhMRs, in indicated conditions (rel. = relative). **(G)** Gene-concept network maps from GO analysis for each indicated cluster. BP = biological process; MF = molecular function; CC = cellular components. Adjusted *p*-value cut-off = 0.05 from GO Enrichment Analysis after Benjamini–Hochberg correction. **(H)** Lists of representative genes associated to DhMRs in each cluster. **(I)** Euler plot showing the overlap of identified DhMRs in eBED or cHFD with respect to the chow condition. **(J)** Representative DhMRs showing eBED-induced increase of 5hmC signal over intron 4 of *Dlgap2* and nearby intronic and exonic regions. **(K)** Representative DhMRs showing eBED-induced decrease of 5hmC signal over intron 1 of *Hmcn1* and nearby intronic and exonic regions. In all panels, DhMRs’ effect size cut-off = 0.05; *p*-value cut-off = 0.01 from Wald statistical testing. *N* = 12 (2 samples each condition – eBED, cHFD, chow – and each mark – 5mC, 5hmC).

Intriguingly, eBED animals showed >900 differentially methylated regions (DMRs; **Supplementary Figure 2B**) and ∈1400 differentially *hydroxy*methylated regions (DhMRs) relative to cHFD controls (**Figure 2B**). The high numbers were surprising because the primary difference between eBED and control animals is *timing* of HFD ingestion, and only for five days. 5mC and 5hmC changes were highly correlated, suggesting that eBED rewires active and regulatory chromatin states (as opposed to unmarked/naïve regions)(**Figure 2C, Supplementary Figure 2C** and **2D**). Given this strong correlation and high absolute levels of 5hmC at DhMRs (**Supplementary Figure 2D**), we focused our analysis on 5hmC changes. eBED-triggered DhMRs were significantly depleted from intergenic regions and CpG islands, and enriched at gene promoters, promoter proximal regions, and over gene bodies (**Figure 2D**). Comparison with ChromHMM annotations showed that DhMRs occured predominantly at strong and permissive transcriptional states (**Figure 2E**). These data are consistent with hydroxymethylation being a dynamic regulatory mark in neurons.^48^ They demonstrate that eBED rapidly and preferentially rewires DNA methylation states over the active cis-regulatory genome. Thus, eBED rapidly rewires the VTA^DA^ epigenome.

Interestingly, eBED-induced 5hmC changes were not correlated with those triggered by the control diet (ad libitum HFD and chow; cHFD) compared to standard chow-only controls (**Figure 2F–2I**). This finding indicated that eBED induces a distinct and highly specific form of dopaminergic epigenome remodeling relative to classic hypercaloric/HFD feeding (**Figure 2I**). Functional Gene Ontology (GO) analysis of eBED-triggered DhMRs identified pathways related to neurodevelopment, synaptic organization, and plasticity, including up-regulation at genes governing neuronal projection, axon guidance, and dopamine signaling, and down-regulation at genes involved in metabolic and cytoskeletal maintenance programs (**Figure 2G,2H, 2J** and **2K**). Thus, eBED causes rapid, specific, plasticity-related epigenome remodeling in VTA^DA^ neurons.

### The hydroxymethylase *Tet1* is enriched in VTA^DA^ neurons

The rapid and targeted nature of the eBED response suggested that one or more TET enzymes might mediate this process. The TET enzymes (TET1-3) oxidize methylated cytosine (5mC) to hydroxymethylcytosine (5hmC), thereby enabling transcriptional activation.^37,49,50^ While all three *Tet* genes are detectably expressed in the VTA, *Tet1* expression is substantially higher than *Tet2* and *Tet3* (**Figure 3A**). Morphologically, we observed that mouse VTA^DA^ neurons exhibit uniquely large, 5hmC-rich nuclei, consistent with a nuclear composition amenable to rapid and widespread epigenetic regulation (**Figure 3B** and **3C**). This distinctive architecture was mirrored (or even exaggerated) in human post-mortem midbrain samples showing conservation of these features (**Figure 3D** and **3E**). Importantly, analysis of *Tet1*^*lacZ*^ knock-in reporter mice (Tet1^tm1Koh^ line^51^) revealed TET1 expression to be restricted to only a few brain regions, including the VTA^DA^ (**Figure 3F**). These results were validated by immunofluorescence (**Figure 3G**) and confirmed robust TET1 (but not TET2) enrichment in the VTA^DA^ compartment. Thus, *Tet1* is enriched in VTA^DA^ neurons.

**Figure 3.**
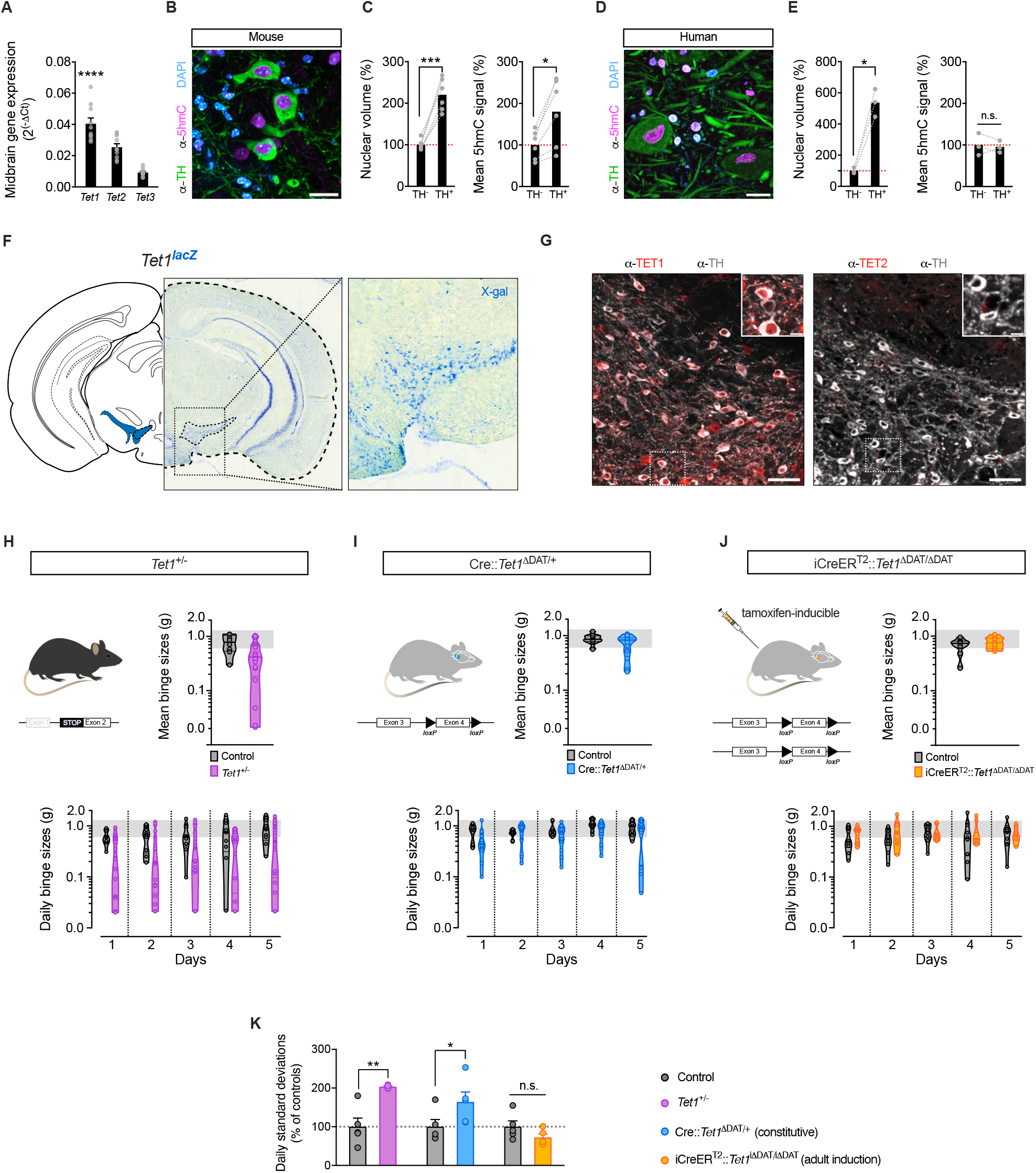
Heterozygous loss of *Tet1* triggers heterogeneity in binge-eating susceptibility. **(A)** qPCR analysis of mRNA expression in midbrain lysates (ΔCt). Data are presented as mean ± SEM. **** *P <* 0.0001. *n* = 10 mice (one-way ANOVA). **(B)** Confocal micrograph of mouse VTA^DA^ neurons (TH^+^; green) immunoreactive for 5hmC (magenta) and nuclear counterstain (DAPI; cyan). Scale bar 20 ţm. **(C)** Nuclear volume and 5hmC signal (mean fluorescence intensity) of murine VTA^DA^ neurons (TH^+^) normalized to TH^−^ cells. Data are presented as mean ± SEM. * *P <* 0.05, ** *P <* 0.01. *n* = 6 mice (paired Student’s *t* -test). **(D)** Confocal micrograph of human VTA^DA^ neurons (TH^+^; green) immunoreactive for 5hmC (magenta) and nuclear counterstain (DAPI; cyan). Scale bar 20 ţm. **(E)** Nuclear volume and 5hmC signal (mean fluorescence intensity) of human VTA^DA^ neurons (TH^+^) normalized to TH^−^ cells. Data are presented as mean ± SEM. * *P <* 0.05. *n* = 3 individuals (paired Student’s *t* -test). **(F)** *Tet1*^*lacZ*^ reporter mouse after X-gal staining; hemisection overview and magnified VTA (insert). **(G)** Confocal micrographs of VTA^DA^ neurons (TH^+^; grey) immunoreactive for TET1 (red) but not TET2; scale bar 50 ţm and 10 ţm (insert). **(H)** *Tet1*^+/−^ mouse model of global *Tet1* haploinsufficiency (left panel). Mean binge sizes (right panel), and daily binge sizes (lower panel) both plotted on a Log_10_ scale (shaded area: normal binge response). Data are presented as violin plots with median and quartiles. *n* = 8–20 mice. **(I)** Cre::*Tet1*^ΔDAT/+^ mouse model of DA neuron-specific *Tet1* heterozygous loss (left panel). Mean binge sizes (right panel), and daily binge sizes (lower panel) both plotted on a Log_10_ scale (shaded area: normal binge response). Data are presented as violin plots with median and quartiles. *n* = 9–17 mice. **(J)** iCreER^T2^::*Tet1*^ΔDAT/ΔDAT^ mouse model for tamoxifen-inducible, DA neuron-specific *Tet1* homozygous loss with adult onset (left panel). Mean binge sizes (right panel) and daily binge sizes (lower panel) both plotted on a Log_10_ scale (shaded area: normal binge response). Data are presented as violin plots with median and quartiles. *n* = 7–10 mice. **(K)** Standard deviation of Log_10_-transformed daily binge sizes relative to respective controls. Data are presented as mean ± SEM of STDEV from Day 1–5. * *P <* 0.05, ** *P <* 0.01 (unpaired Student’s *t* -test).

### *Tet1-*haploinsufficiency triggers marked inter-individual variation in eBED susceptibility

Given the prominent hydroxymethylome changes and *Tet1* enrichment in VTA^DA^ neurons, we asked whether *Tet1* loss might alter susceptibility to binge eating. We used the same *Tet1*^*lacZ*^ reporter line, in which full-length *Tet1* is disrupted during development (Tet1^tm1Koh^; Khoueiry et al., 2017^51^). Because homozygous deletion causes embryonic abnormalities, we focused on heterozygous mice, thus modelling both partial gene-dosage effects and a range of variation expected in human populations.^52^ *Tet1*^*+/–*^ mice were born at Mendelian ratios, appeared normal, and showed no differences in chronic HFD-induced weight gain, food intake, food preference, energy expenditure, respiratory exchange ratio (RER), or voluntary wheel running activity when compared to littermate controls (**Supplementary Figure 3A**). Thus, *Tet1* haploinsufficiency does not alter energy homeostasis, overall feeding behavior, or dietary preference.

Intriguingly, however, *Tet1*^*+/–*^ mice displayed pronounced inter-individual heterogeneity when challenged with eBED. Specifically, within any given *Tet1*^*+/–*^ isogenic cohort, we observed subsets of heterozygotes that consistently exhibited *resilience* to eBED, and others that consistently showed high *susceptibility* (**Figure 3H**). This hypervariable phenotype was absent in wild-type littermates and was highly reproducible across litters, breeding pairs, and observed in both males and females (**Supplementary Figure 3B**). The effect was specific to eBED, as *Tet1*^*+/–*^ mice did not show increased variability in anxiety-like behavior (elevated plus maze; **Supplementary Figure 3C**) or in cocaine-conditioned place preference (**Supplementary Figure 3D**). Thus, *Tet1* haploinsufficency triggers marked heterogeneity in inter-individual binge-eating susceptibility.

Similar ‘prone’ and ‘resilient’ intra-strain states have been observed in other food addiction contexts^53–55^ and our data suggested that *Tet1* could lie mechanistically upstream of such individuality. To better understand the phenotype and to confirm specificity, we generated two additional loss-of-function models. Importantly, conditional deletion of *Tet1* in dopaminergic neurons (*DAT-Cre* x *Tet1*^flox/+^; referred to as Cre::*Tet1*^Δ*DAT/+*^ mice) produced conditional heterozygous mutants that also showed exaggerated inter-individual variability in binge-eating behavior (**Figure 3I** and **Supplementary Figure 3E**), albeit less dramatic. This finding validated the phenotype and localized the origin of the effect to dopaminergic neurons. Moreover, we tested adult-onset, dopamine neuron-specific deletion of both copies of *Tet1* using a tamoxifen-inducible *DAT-iCreER*^*T2*^ system. Intriguingly, adult-specific *Tet1* deletion mutants (iCreER^T2^::*Tet1*^ΔDAT/ΔDAT^ mice) showed no evidence of an altered variability phenotype (**Figure 3J** and **Supplementary Figure 3F**). Because both *Tet1*^*+/–*^ and Cre::*Tet1*^Δ*DAT/+*^ disrupt *Tet1* during development, and since adult-onset deletion did not alter variability, these data suggest that *Tet1* dosage influences susceptibility through an early-life mechanism. Thus, developmental dopaminergic *Tet1* dosage regulates binge-eating susceptibility.

### *Tet1* shapes VTA input architecture

Because developmental, but not adult-induced, *Tet1* deletion triggered eBED heterogeneity, we hypothesized that *Tet1* dosage influences the establishment of VTA^DA^ circuitry. Consistent with this idea, *Tet1* is expressed at high levels in the embryonic ventral mesencephalon (**Supplementary Figure 4A** and **4B**) during the period when VTA^DA^ neurons are generated^56,57^ and remains highly expressed in developing VTA^DA^ neurons^58^ (**Supplementary Figure 4C**). This developmental period overlaps with the establishment of the VTA’s input architecture from sensory, limbic and cortical regions that shape VTA^DA^ responsiveness. Notably, immunofluorescence analyses showed that dopamine neuron numbers across VTA subregions were unchanged in adult *Tet1*^*+/–*^ mice, regardless of whether they were eBED-prone or eBED-resilient (**Supplementary Figure 4D** and **4E**). Thus, *Tet1* dosage does not alter overall dopaminergic cell number or gross VTA organization.

To test whether dosage instead influenced circuit topology and/or input balance, we applied virus-assisted retrograde tracing to *Tet1*^*+/–*^ littermates (**Figure 4A, Supplementary Figure 4F**). We injected a retrograde AAV variant with preferential uptake at nerve terminals (AAVrg-hSyn1-eGFP^59^), into the VTA of 8 w-old adult males, allowed four weeks for retrograde transport and recovery, phenotyped each animal for eBED responsiveness, and quantified labeled projection neurons by eGFP-fluorescence. Retrogradely labelling were found in the expected input regions,^17,60,61^ including the lateral hypothalamic area (LHA), the dorsal and ventral striatum (d/vSTR), the nucleus accumbens (NAc), the ventral pallidum (VP), and the medial prefrontal cortex (mPFC), including the prelimbic (mPFC^PL^) and anterior cingulate (mPFC^Cg^) regions (**Figure 4B** and **4C**). Intriguingly, binge-resilient *Tet1*^*+/–*^ mice exhibited a significant reduction in back-labelled GFP^+^ neurons in the mPFC^PL^ compared to binge-*prone Tet1*^*+/–*^ littermates (**Figure 4D–4F**). These data demonstrate that *Tet1*^*+/–*^ haploinsufficiency modulates eBED susceptibility in concert with VTA^DA^ input architecture. They suggested that reduced mPFC^PL^→ VTA connectivity might control the interindividual eBED susceptibility thresholds. Thus, *Tet1* dosage controls VTA input architecture.

**Figure 4.**
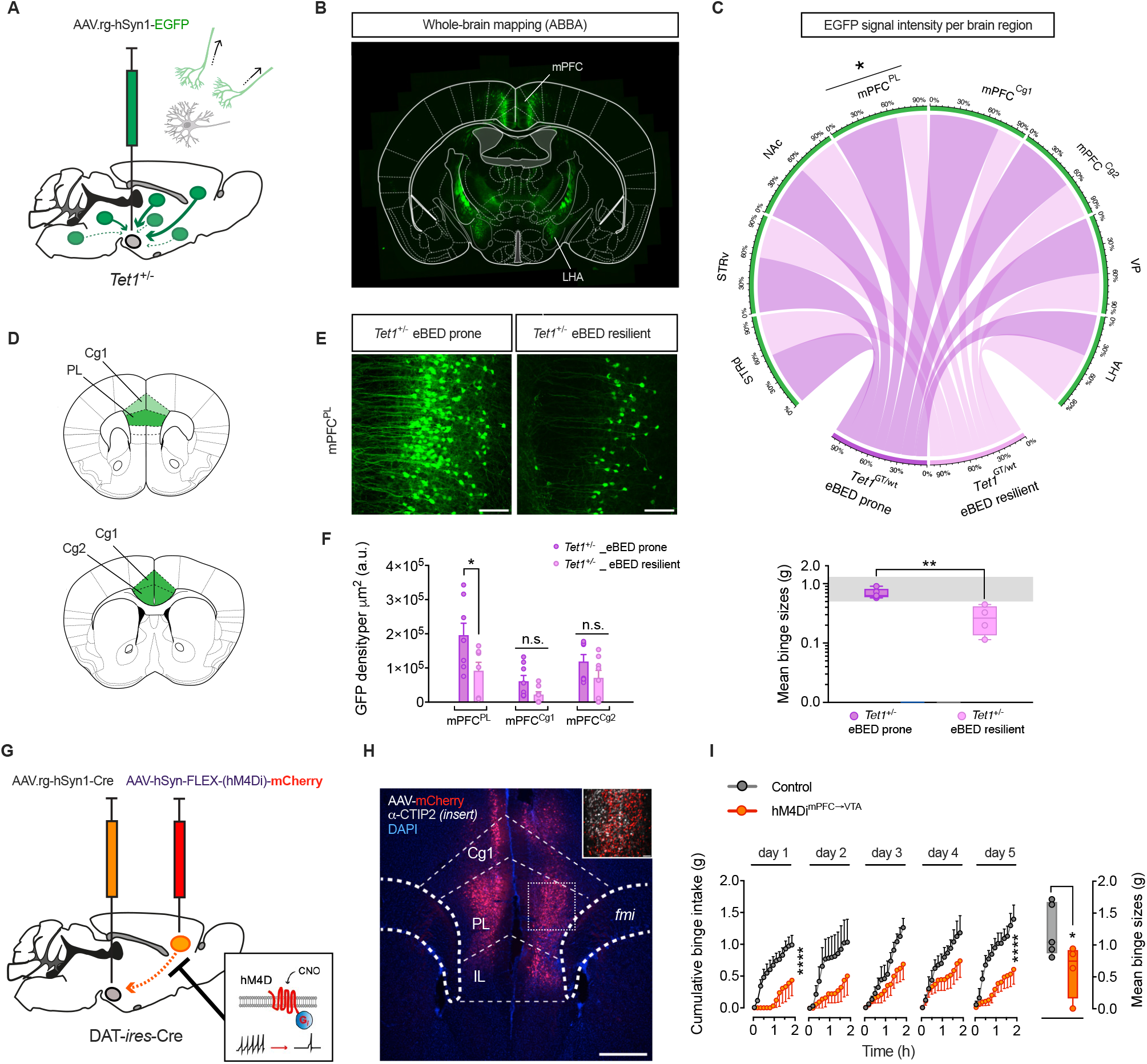
Reduced mPFC^PL^→VTA inputs confer binge-eating resilience in *Tet1*^+/–^ mice. **(A)** Schematic of retrograde labeling of input projection neurons to the VTA using AAV.rg-hSyn1-EGFP. **(B)** Micrograph of an exemplary coronal brain section upon machine learning-assisted brain region mapping (Aligning Big Brains & Atlases (ABBA/ImageJ)). Back-labeled neurons are highlighted in the mPFC and LHA. **(C)** Circos plot showing relative input strength from reward-related regions to the VTA comparing eBED-prone versus -resilient *Tet1*^+/–^ mice (upper panel); corresponding binge sizes of individual *Tet1*^*+/-*^ mice across 5 Ebed days plotted on a Log_10_ scale to visualize varied responses (lower panel; shaded area shows normal binge response). Data are presented as mean minimum to maximum values. ** *P <* 0.01. *n* = 4 mice (unpaired Student’s *t* -test). **(D)** Coronal brain schematics of mPFC subregions (*PL*: prelimbic; *Cg1*: dorsal cingulate cortex; *Cg2*: ventral cingulate cortex). **(E)** Representative micrographs of mPFC^PL^ from eBED-prone versus -resilient *Tet1*^+/−^ mice. Scale bar 100 ţm. **(F)** Bar graphs of raw EGFP density values from mPFC subregions hemisections (*PL*: prelimbic; *Cg1*: dorsal cingulate cortex; *Cg2*: ventral cingulate cortex). Data are presented as mean ± SEM. * *P <* 0.05. *n* = 4 mice (two-tailed unpaired Student’s *t* -test). **(G)** Schematic illustrating chemogenetic approach to selectively inhibit VTA-projecting mPFC^PL^ neurons using a dual AAV approach (AAV.rg-Cre + AAV-hSyn1-FLEX-hM4Di-mCherry). **(H)** Micrograph of VTA-projecting mPFC^PL^ neurons expressing mCherry (red) intermingled with layer-5 pyramidal neurons (CTIP2: grey). Scale bar: 500 ţm and 50 ţm (insert). **(I)** Cumulative binge intake of hM4Di^mPFC-VTA^ mice and controls per individual days (left panel). Data are presented as mean ± SEM. *** *P <* 0.001, **** *P <* 0.0001. *n* = 4–5 mice (multiple unpaired Student’s *t* -test); mean binge sizes of hM4Di^mPFC-VTA^ mice and controls (right panel). Data are presented as mean and minimum to maximum. * *P <* 0.05, *n* = 4–5 mice (one-tailed unpaired Student’s *t* -test).

### Blunted mPFC^PL^VTA activity induces eBED resilience

To test whether the altered mPFC^PL^→VTA connectivity might underpin the observed eBED susceptibility differences, we used a dual-AAV chemogenetic approach, combining retrograde labelling with chemogenetic inhibition. We injected AAVrg-hSyn1-*Cre* into the VTA of mice to functionally back-label input neurons with *Cre* recombinase, and AAV-FLEX-hM4Di-mCherry into the mPFC^PL^, thereby restricting expression of the inhibitory hM4Di receptor to VTA-projecting mPFC^PL^ neurons (**Figure 4G**). Correct targeting was confirmed by mCherry and CTIP2 immunoreactivity in layer 5 pyramidal neurons (**Figure 4H**). Following recovery, mice were exposed to 4 weeks of ad libitum HFD and chow, followed by two days of HFD withdrawal, and then subjected to the five-day eBED paradigm. All animals received CNO 15 min prior to each 2 h-HFD binge opportunity to acutely silence mPFC^PL^→VTA projections. Critically, whereas control mice (expressing mCherry alone) displayed the expected escalation in binge size across days, those subject to chemogenetic inhibition showed an abolished response (**Figure 4I**), and tended to exhibit reduced meal sizes and eating rate (**Supplementary Figure 4G** and **4H**). Cumulative chow intake was not different between groups, arguing against influences of divergent energy balance (**Supplementary Figure 4I**). These data demonstrated that acute inhibition of mPFC^PL^→VTA activity is sufficient to cause eBED resilience. Thus, blunted mPFC^PL^→VTA^DA^ activity decreases binge-eating susceptibility.

### EGR1-guided TET1 reactivation restores eBED susceptibility

Collectively, the data above identified *Tet1* dosage as a determinant of inter-individual binge-eating susceptibility and linked the observation to control of VTA^DA^ circuit connectivity. We next asked whether this *Tet1*-dependent axis remains accessible in adulthood. Motif enrichment analysis of the eBED-induced VTA^DA^ 5hmC response (DhMRs) revealed prominent binding motifs for general transcriptional regulators such as TATA-box binding protein (TBP) and select zinc-finger motifs (e.g. ZNF281), but also an enrichment for Early Growth Response 1 (EGR1)(**Figure 5A–5C** and **Supplementary Figure 5A**), a rapidly inducible transcription factor known to recruit TET1, and to act in dopaminergic neurons.^62–66^ Immunofluorescence confirmed robust EGR1 protein induction in VTA^DA^ neurons upon eBED (**Figure 5D** and **5E**). Consistent with its activity-dependent nature, the number of EGR1^+^ cells increased progressively from 1 to 4 h upon eBED (**Figure 5F**). Notably, this induction appeared specific to the VTA; substantia nigra (SNc) neurons showed no significant change (**Figure 5G**). Also noteworthy, c-Fos^+^ neuron counts (another activity-induced transcription factor) were stable across conditions (**Supplementary Figure 5B**), indicating that EGR1 activation represents a specific molecular signature of the binge-eating response. Thus, eBED triggers acute induction of EGR1.

**Figure 5.**
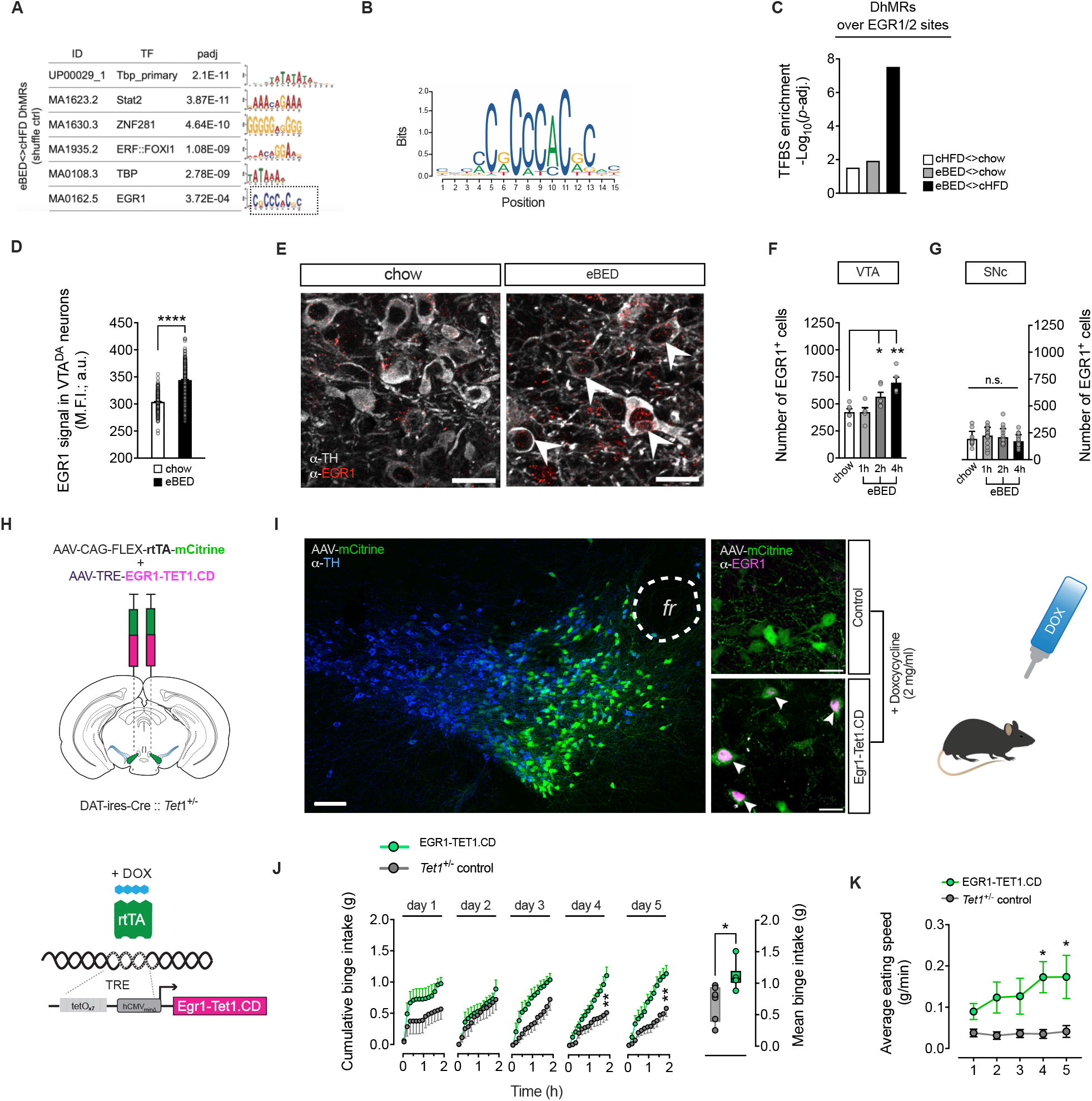
EGR1-guided TET1 reactivation in VTA^DA^ neurons restores binge-eating susceptibility. **(A)** Motif analyses on eBED-induced DhMRs showing significant enrichment of indicated transcription factors, including EGR1. Adjusted *p*-values from one-tailed Fisher’s exact test, followed by Bonferroni correction. **(B)** Sequence logo representation of the EGR1/2 binding motif from JASPAR database, used for enrichment analysis in C. **(C)** Bar plots showing the enrichment of EGR1/2 binding sites at DhMRs, across indicated comparisons. Adjusted *p*-values from one-tailed Fisher’s exact test, followed by Bonferroni correction. **(D)** Quantification of EGR1 signal (mean fluorescence intensity; M.F.I.) in VTA^DA^ neurons at 2 h after eBED versus chow fed control mice. Data are presented as mean ± SEM. **** *P <* 0.0001. *n* = 4–7 mice (unpaired Student’s *t* -test). **(E)** Representative micrographs of EGR1 immunoreactivity (red) co-localizing to VTA^DA^ neurons (TH^+^, grey). Scale bars: 20 ţm. **(F)** Quantification of EGR1 immunoreactivity punctae in the VTA at 1 h, 2 h or 4 h after eBED versus chow-fed control mice. Data are presented as mean ± SEM. * *P <* 0.05, ** *P <* 0.01. *n* = 4–7 mice (one-way ANOVA). **(G)** Quantification of EGR1 immunoreactivity punctae in the SNc at 1 h, 2 h or 4 h after eBED versus chow fed control mice. Data are presented as mean ± SEM. * *P <* 0.05, ** *P <* 0.01. *n* = 4–7 mice (one-way ANOVA). **(H)** Schematic depicting experimental approach for viral targeting of VTA^DA^ neurons for doxycycline-inducible expression of an EGR1-TET1.CD fusion construct in *Tet1*^+/−^ mice. **(I)** Viral targeting as confirmed by mCitrine viral expression (green) co-localizing with VTA^DA^ neurons (TH^+^, blue). EGR1 immunoreactivity (magenta) was robustly expressed in VTA^DA^ mCitrine^+^ neurons (green) from EGR1-TET1.CD, but not from *Tet1*^+/–^ control mice. Scale bar: 50 ţm and 10 ţm (middle panels). **(J)** Cumulative binge intake of EGR1-TET1.CD mice and *Tet1*^+/−^ controls per individual days (left panel). Data are presented as mean ± SEM. ** *P <* 0.01. *n* = 6 mice (multiple unpaired Student’s *t* -test). Summary data of mean binge sizes (right panel). Data are presented as mean ± SEM. * *P <* 0.05. *n* = 6 mice (one-tailed unpaired Student’s *t* -test; right panel). **(K)** Average eating speed of EGR1-TET1.CD mice and *Tet1*^+/−^ controls per individual days. Data are presented as mean ± SEM. * *P <* 0.05, *n* = 6 mice (two-way ANOVA).

EGR1 has previously been shown to recruit TET1 to its genomic binding sites and to regulate activity-dependent transcription in the brain.^64^ Based on the enrichment and induction of EGR1 observed during eBED, we fused full-length EGR1 to the catalytic domain of TET1^67^ (EGR1-TET1.CD), using previously established structural constraints^64^ (**Supplementary Figure 5C**). We then used this fusion protein to actively target TET1 activity to EGR1 binding sites in vivo. To mimic the acute induction observed physiologically, we used a dual-AAV system combining Cre-dependent expression of rtTA in VTA^DA^ neurons with doxycycline (DOX)-inducible transgene activation (**Figure 5H**).

Adult *Tet1*^*+/–*^ mice received either both AAVs to enable inducible expression of the EGR1-TET1.CD fusion protein in VTA^DA^ neurons (EGR1-Tet1.CD mice) or a single control AAV (*Tet1*^*+/–*^ controls). After recovery, mice were acclimated to HFD (+chow) for four weeks followed by a two-day chow-only withdrawal. After five days of doxycycline treatment, robust EGR1 immunoreactivity was detected selectively in VTA^DA^ neurons of EGR1-Tet1.CD mice, confirming precise induction and spatial specificity of the system (**Figure 5I**). DOX induction was initiated on Day 1 of the eBED paradigm. As expected, a subset of *Tet1*^*+/–*^ control mice remained resilient to binge-like behavior. In striking contrast, all EGR1-Tet1.CD mice showed robust binge eating, with significantly elevated binge size and eating speed (**Figure 5J** and **5K**). Thus, targeted reinstatement of TET1 catalytic activity restores binge-eating susceptibility across *Tet1*^+/–^ mice. These findings show that binge-eating susceptibility setpoints are accessible to *Tet1*-dependent mechanisms in adulthood. The findings collectively identify Tet1 as a central regulator of inter-individual variation in binge-eating susceptibility.

### *TET1* links (epi)genetic variation, reward-circuit function and binge eating in humans

To test whether this *Tet1*-dependent axis might extend to humans, we examined the relationship between *TET1* promoter methylation, reward-circuit function (BOLD fMRI), and binge-eating behavior in ESTRA^68^ and IMAGEN,^69^ cohorts comprising individuals with eating disorders and age- and sex-matched controls (*n* = 145; all female; **Figure 6A**). We first selected CpG sites within the TET1 promoter (**Figure 6B**) and evaluated their suitability as peripheral surrogates of brain epigenetic variation by examining blood–brain methylation correspondence using the Blood–Brain DNA Methylation Comparison Tool.^70^ Among these sites, cg23602092 showed strong cross-tissue correlation, particularly in the prefrontal cortex (PFC; *r*^2^ = 0.77, *P* = 3.98 × 10^−16^; **Table 1**). Intriguingly, higher methylation at cg23602092 was significantly associated with increased binge-eating frequency in ESTRA (*β* = 0.0145, *P* = 0.014; **Figure 6C**). Consistent with prior reports,^71^ cg23602092 displayed a bimodal distribution (**Figure 6D**), separating participants into high- and low-methylation subgroups (*n* = 17 and 129, respectively). Individuals in the high-methylation subgroup exhibited significantly greater binge-eating frequency (*P* = 0.042; **Figure 6E**). Thus, methylation at cg23602092 marks an epigenetic state associated with increased binge-eating susceptibility in humans.

**Table 1.** Correlations between blood and brain DNA methylation levels at CpG sites within the *TET1* promoter region.

| Blood Brain DNA Methylation Comparison<br>[Bonferroni corrected $p$-value threshold = $0.05/(24 \times 4) = 5.21 \times 10^{-4}$] | | | | | | | | | | | |
| --- | --- | --- | --- | --- | --- | --- | --- | --- | --- | --- | --- |
| CpG | Position hg38 |  |  | Prefrontal cortex |  | Entorhinal cortex |  | Superior temp. gyrus |  | Cerebellum |  |
| | chr | start | end | $r$ | $p$ | $r$ | $p$ | $r$ | $p$ | $r$ | $p$ |
| cg15538753 | chr10 | 68 558 238 | 68 558 239 | 0.0975 | 0.408 | 0.0317 | 0.793 | -0.1350 | 0.249 | 0.0954 | 0.429 |
| cg23602092 | chr10 | 68 559 888 | 68 559 889 | 0.7700 | 3.980E-16 | 0.6810 | 6.310E-11 | 0.7420 | 2.660E-14 | 0.5500 | 6.650E-7 |
| cg22876739 | chr10 | 68 560 017 | 68 560 018 | -0.0400 | 0.721 | -0.0960 | 0.428 | 0.0540 | 0.648 | 0.0610 | 0.612 |
| cg02952701 | chr10 | 68 560 026 | 68 560 027 | -0.0880 | 0.456 | 0.0115 | 0.924 | -0.0608 | 0.604 | -0.0352 | 0.771 |
| cg00128195 | chr10 | 68 560 241 | 68 560 242 | 0.3990 | 4.240E-4 | 0.4260 | 2.140E-4 | 0.2170 | 0.062 | 0.1100 | 0.359 |
| cg07669489 | chr10 | 68 560 280 | 68 560 281 | 0.0601 | 0.611 | 0.0854 | 0.479 | 0.3680 | 1.170E-3 | 0.0293 | 0.808 |
| cg10413954 | chr10 | 68 560 282 | 68 560 283 | -0.0212 | 0.858 | -0.1110 | 0.357 | 0.1090 | 0.353 | 0.0502 | 0.677 |
| cg21948169 | chr10 | 68 560 294 | 68 560 295 | 0.0011 | 0.993 | 0.0821 | 0.496 | -0.0088 | 0.940 | 0.1190 | 0.323 |
| cg05400741 | chr10 | 68 560 316 | 68 560 317 | 0.1400 | 0.235 | 0.1750 | 0.145 | 0.0834 | 0.477 | 0.0369 | 0.760 |
| cg03651138 | chr10 | 68 560 456 | 68 560 457 | 0.1860 | 0.112 | 0.2330 | 0.051 | 0.1040 | 0.377 | -0.0092 | 0.939 |
| cg19439331 | chr10 | 68 560 713 | 68 560 714 | 0.1220 | 0.299 | -0.0328 | 0.786 | 0.0503 | 0.668 | 0.1510 | 0.209 |
| cg03756448 | chr10 | 68 560 775 | 68 560 776 | 0.0663 | 0.574 | 0.0223 | 0.853 | 0.0967 | 0.409 | 0.0795 | 0.510 |
| cg09183181 | chr10 | 68 560 977 | 68 560 978 | -0.0131 | 0.912 | 0.0435 | 0.719 | 0.0739 | 0.529 | 0.0013 | 0.992 |
| cg13848707 | chr10 | 68 561 486 | 68 561 487 | 0.1470 | 0.211 | -0.3110 | 8.220E-3 | 0.1380 | 0.238 | -0.0257 | 0.832 |
| cg06767766 | chr10 | 68 561 626 | 68 561 627 | -0.0484 | 0.682 | 0.1390 | 0.247 | -0.0185 | 0.875 | -0.0878 | 0.466 |
| cg23350336 | chr10 | 68 561 797 | 68 561 798 | 0.2360 | 0.043 | 0.0263 | 0.828 | 0.0922 | 0.432 | -0.0089 | 0.941 |
| cg12630147 | chr10 | 68 561 817 | 68 561 818 | 0.1530 | 0.193 | 0.1010 | 0.403 | 0.0327 | 0.781 | 0.1020 | 0.396 |
| cg02774862 | chr10 | 68 561 823 | 68 561 824 | 0.1620 | 0.167 | -0.1630 | 0.175 | 0.1050 | 0.369 | -0.1050 | 0.383 |
| cg09978996 | chr10 | 68 561 911 | 68 561 912 | -0.1100 | 0.353 | 0.0121 | 0.920 | 0.1980 | 0.089 | -0.1480 | 0.218 |
| cg15254238 | chr10 | 68 562 117 | 68 562 118 | 0.1400 | 0.233 | -0.0326 | 0.787 | 0.2540 | 0.028 | -0.9180 | 0.446 |
| cg25926515 | chr10 | 68 562 132 | 68 562 133 | 0.0708 | 0.549 | 0.1420 | 0.238 | 0.1830 | 0.117 | -0.0964 | 0.424 |
| cg10403849 | chr10 | 68 562 202 | 68 562 203 | 0.0407 | 0.731 | -0.0747 | 0.536 | -0.1230 | 0.295 | -0.2280 | 0.056 |
| cg19127638 | chr10 | 68 562 685 | 68 562 686 | 0.2250 | 0.053 | 0.2420 | 0.042 | 0.2770 | 0.016 | 0.1600 | 0.182 |
| cg17817532 | chr10 | 68 563 117 | 68 563 118 | 0.1250 | 0.289 | 0.0951 | 0.430 | 0.0616 | 0.599 | 0.0107 | 0.929 |

**Figure 6.**
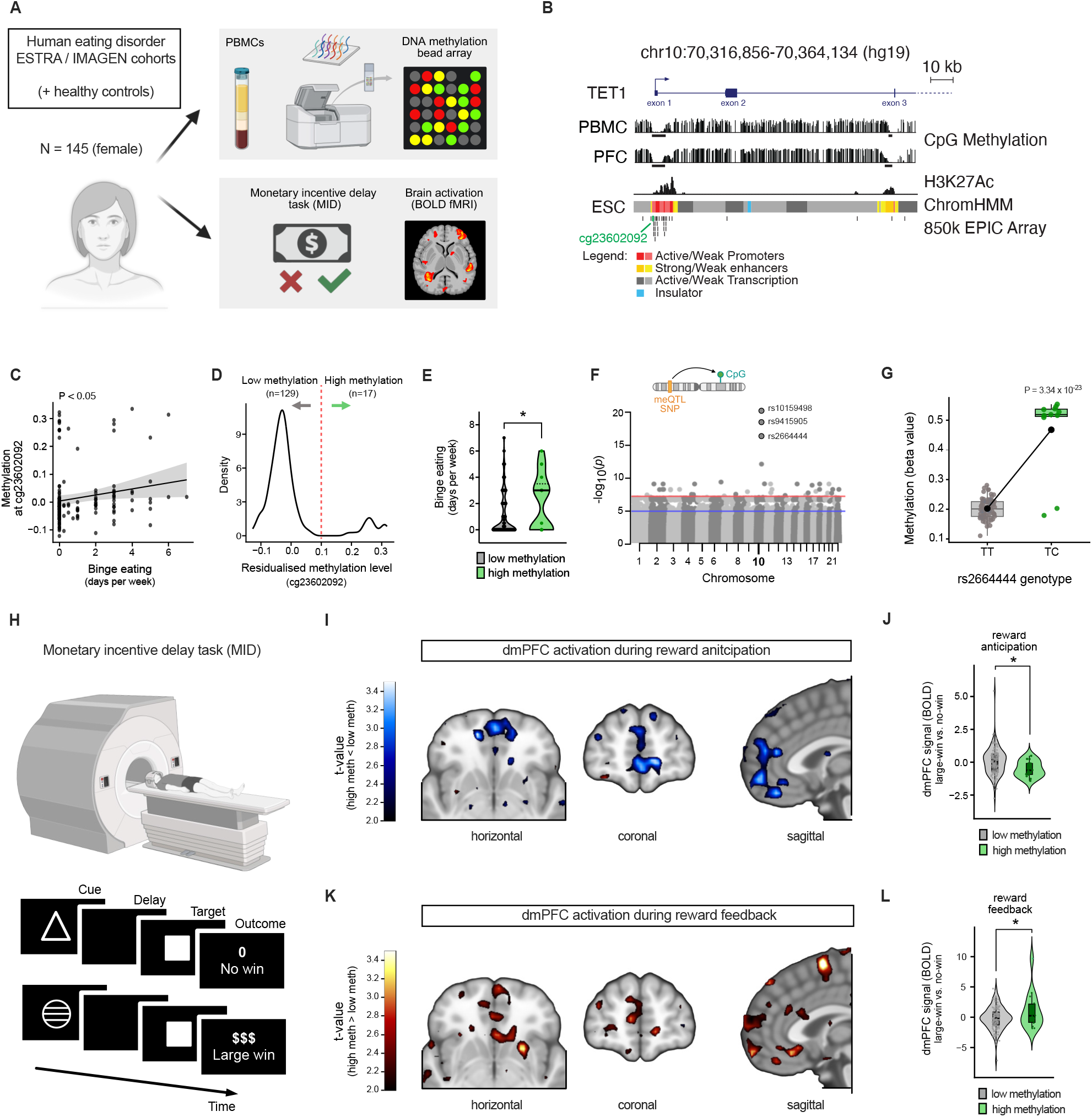
Human *TET1* methylation status mediates binge eating and dmPFC brain activation during reward processing. **(A)** Individuals with and without eating disorders from the ESTRA^68^ and IMAGEN^69^ cohorts were screened for their PBMC methylome (Illumina Human Methylation EPIC v2.0 array; upper panel); the same individuals additionally underwent the monetary incentive delay task (MID) to assess reward-circuit function using BOLD fMRI (lower panel). *Created in BioRender.com*. **(B)** Genomic snapshot of the *TET1* human gene promoter and first 3 exons, from the hg19 genome reference in the UCSC genome browser. DNA methylation levels of peripheral blood mononuclear cells (PBMC) and prefrontal cortex cells (PFC) are shown from Methbase. Solid black bars below DNA methylation tracks represent hypomethylated regions. The H3K27Ac track shows the solid overlay of signals from seven cell lines from ENCODE (GM12878, H1-hESC, HSMM, HUVEC, K562, NHEK, NHLF). Exon 1, a non-transcribed regulatory region, showing congruent hypomethylation and H3K27ac-enrichment in both PBMC and PFC samples. Chromatin region classification is from ChromHMM pipeline on H1-human ESCs. The probes from the human 850k EPIC Array on CpG sites are reported. Cg23602092 (indicated in green), was found to be differentially methylated and segregating individuals into either low or high methylation subgroups. **(C)** Linear regression analysis showing a significant positive relationship between methylation status at cg23602092 and binge-eating frequency. **(D)** Density distribution of cg23602092 methylation across all subjects revealing a bimodal pattern separating individuals into low- and high methylation subgroups (*n* = 129 and 17, respectively; cut-off at 0.1 residualised methylation level). **(E)** Quantification of binge-eating frequency in low methylation and high methylation subgroups. Data are presented as violin plot with median and quartiles. * *P <* 0.05. *n* = 129 and 17, respectively (unpaired Student’s *t* -test). **(F)** Manhattan plot showing three cis-meQTLs on chromosome 10. **(G)** Representative cis-meQTL rs2664444_C (minor allele) is strongly associated with cg23602092 methylation. **(H)** Illustration of the BOLD fMRI paradigm (upper panel) combined with the monetary incentive delay task (MID; lower panel) assessing brain activation during reward anticipation (delay) and reward feedback (outcome); *‘large win’* signal is normalized to *‘no win’* responses. **(I)** dmPFC activation during reward anticipation shown in horizontal, coronal, and sagittal orientation comparing high-versus low methylation groups. **(J)** Corresponding BOLD quantification during reward anticipation (*‘large win’* versus *‘no win’* ). Data are presented as violin-and-box plot with median and quartiles. * *P <* 0.05. *n* = 114 and 14, respectively (unpaired Student’s *t* -test). **(K)** dmPFC activation during reward feedback shown in horizontal, coronal, and sagittal orientation comparing high-versus low methylation groups. **(L)** Corresponding BOLD quantification during reward feedback (*‘large win’* versus *‘no win’* ). Data are presented as violin-and-box plot with median and quartiles. * *P <* 0.05. *n* = 114 and 14, respectively (unpaired Student’s *t* -test).

The bimodal distribution and strong blood–brain concordance also suggested potential underlying genetic regulation. Indeed, cis-meQTL analysis identified four variants (rs10159498, rs9415905, rs2664444, rs1856831) robustly associated with methylation at this site. Importantly, these findings were replicated in an independent adolescent cohort (ARIES)^72^ (**Figure 6F** and **6G**; **Table 2**). Three of these variants, including rs2664444, also showed nominal associations with binge-eating frequency (**Table 3**), thereby linking genetic variation at the *TET1* locus to behavioral outcomes. Participants in the high cg23602092 methylation subgroup were markedly more likely to carry the rs2664444 minor allele (odds ratio = 1371; *P* = 8.69 × 10^−6^), consistent with genetically driven epigenetic stratification. Notably, rs2664444 was also identified as an eQTL for *TET1*, with the minor C allele associated with increased *TET1* expression in the cerebellum (normalized effect size = 0.42; *P* = 1.6 × 10^−4^). The latter finding supports functional relevance of this variant. Thus, genetic variation at the *TET1* locus is associated with coordinated differences in promoter methylation and TET1 expression in humans. We next tested whether this genetically anchored epigenetic variation translated into altered neural processing of reward. Functional MRI data acquired during a reward monetary incentive delay task (**Figure 6H**) revealed significant group differences in activation of the dorsomedial prefrontal cortex (dmPFC), a region homologous to the murine mPFC^PL^ (**Supplementary Figure 6A** and **6B**). Participants with higher cg23602092 methylation (or carriers of the linked rs2664444 C allele) showed reduced activation during reward anticipation compared with lower-methylation individuals or non-carriers (**Figure 6I**). In contrast, the same subgroup exhibited increased dmPFC activation during reward feedback (**Figure 6J**). These data demonstrated that higher *TET1* promoter methylation is associated with altered dmPFC reward processing in humans, paralleling the mPFC→VTA circuitry implicated in *Tet1*^+/–^ mice. Thus, variation at the TET1 locus links epigenetic state, reward-circuit function, and binge-eating behavior in humans.

**Table 2.**
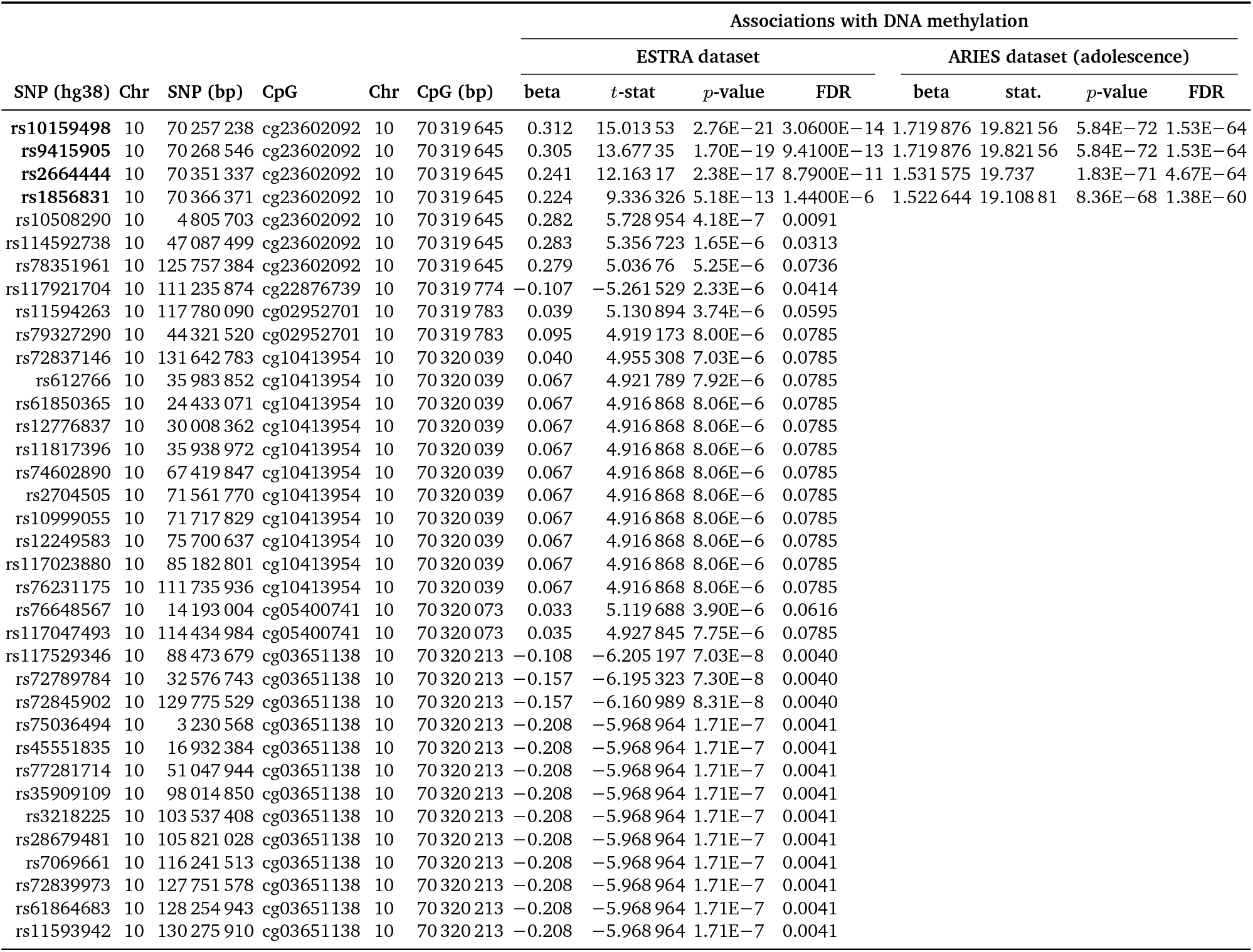
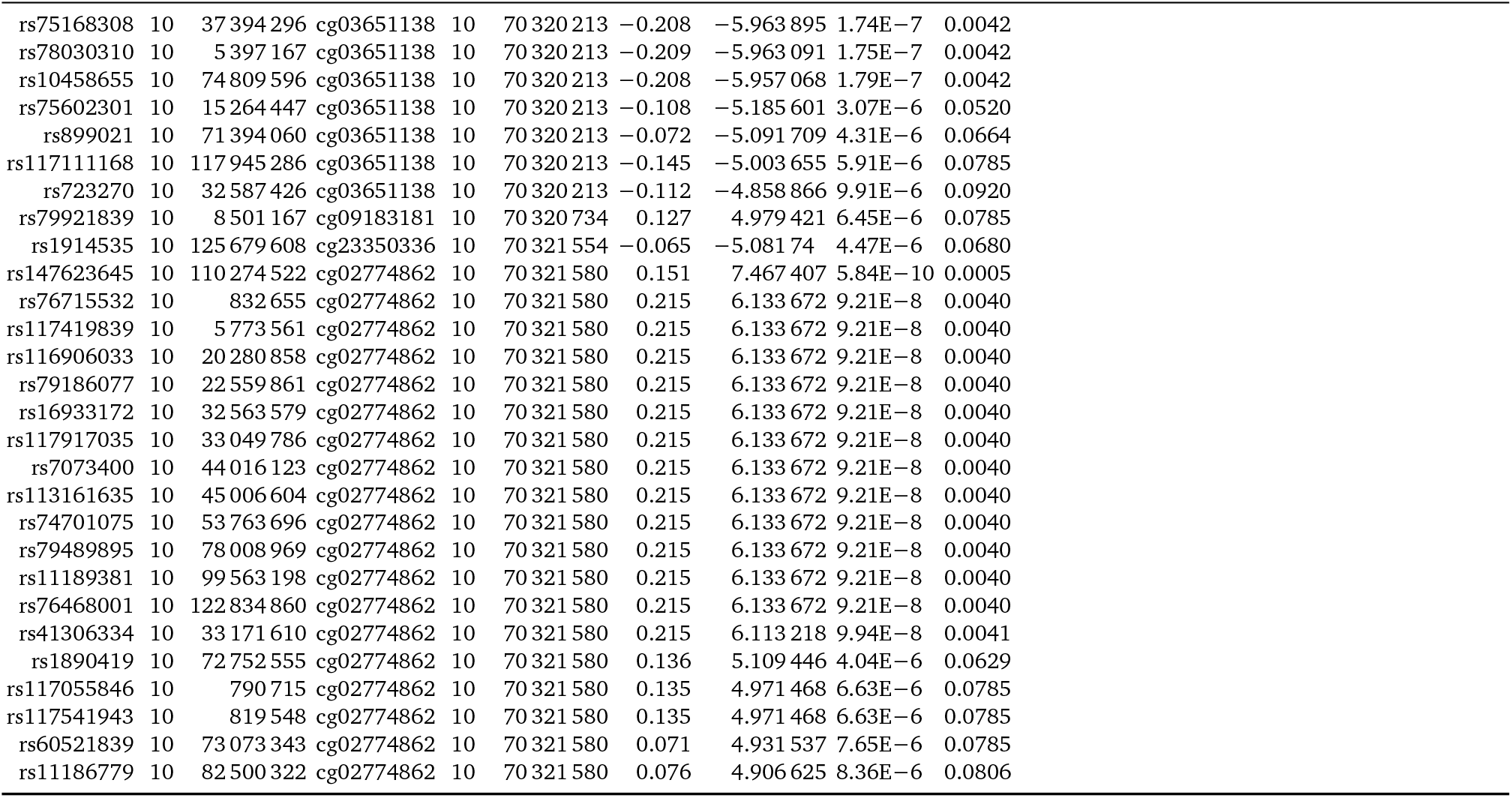
Identification of cis-meQTLs associated with DNA methylation at *TET1* promoter CpGs.

**Table 3.**
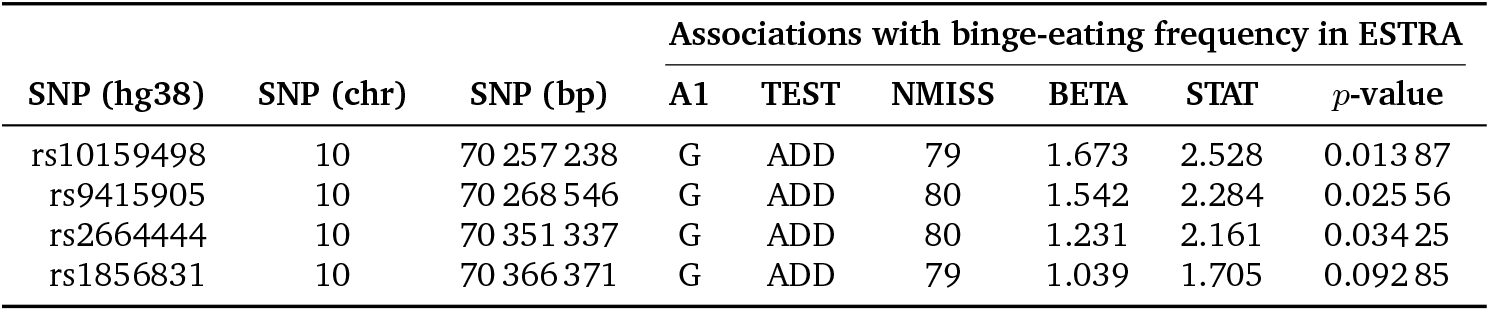
cis-meQTLs associations with binge-eating frequency in ESTRA.

| SNP (hg38) | SNP (chr) | SNP (bp) | Associations with binge-eating frequency in ESTRA |  |  |  |  |  |
| --- | --- | --- | --- | --- | --- | --- | --- | --- |
|  |  |  | A1 | TEST | NMISS | BETA | STAT | p-value |
| rs10159498 | 10 | 70 257 238 | G | ADD | 79 | 1.673 | 2.528 | 0.013 87 |
| rs9415905 | 10 | 70 268 546 | G | ADD | 80 | 1.542 | 2.284 | 0.025 56 |
| rs2664444 | 10 | 70 351 337 | G | ADD | 80 | 1.231 | 2.161 | 0.034 25 |
| rs1856831 | 10 | 70 366 371 | G | ADD | 79 | 1.039 | 1.705 | 0.092 85 |

## Discussion

Evidence from twin studies and from isogenic, environmentally homogeneous animal models indicates that a substantial fraction of behavioral individuality cannot be readily attributed to genetic or environmental variation alone. Monozygotic twins frequently diverge in traits related to motivation, reward seeking, and psychopathology, and genetically identical animals raised under controlled conditions can develop stable, reproducible differences in behavior. Despite substantial research, the molecular mechanisms that generate such individuality have remained enigmatic.^29,73–75^

Here, we identify Tet1 as a regulator of reproducible inter-individual variation that emerges even under shared genetic and environmental conditions. Our data indicate that Tet1 dosage buffers the reproducibility of dopaminergic circuit development such that modest probabilistic divergence in reward-circuit wiring can give rise to meta-stable differences in individual binge-eating susceptibility. When *Tet1* dosage is intact, VTA^DA^ circuitry assembles with high consistency; when dosage is reduced, stochastic variability in input wiring emerges, yielding distinct behavioral phenotypes even among genetically identical individuals. This provides a molecular basis for long-standing observations that behavioral individuality is an intrinsic property of neural systems rather than experimental noise. The findings also provide an example that epigenetic regulation during development can leave lasting *physical* imprints on circuit architecture that may bias future behavioral responses, even in the absence of ongoing molecular perturbation.

Robustness in neural systems reflects the reproducibility of circuit architecture and the stability of cellular function, both of which depend on precisely regulated genetic and epigenetic programs.^76,77^ Neuroanatomical and behavioral variability despite identical genotypes is a widespread phenomenon,^26,27,78–81^ as is epigenetic variation underlying visible traits such as pigmentation and body morphology. Studies have traced some of this variation to metastable epialleles— genomic loci that acquire stable yet variably penetrant methylation or hydroxymethylation states during early development^82,83^—providing a molecular framework for individuality under shared genetic and environmental conditions. Our data extend this concept by showing that modest shifts in gene dosage of an epigenetic regulator are sufficient to modulate variance in behavioral setpoints. To our knowledge, this constitutes identification of one of the first molecular regulators of individuality.

Consistent with this view, TET enzymes are required for normal neurodevelopment and transcriptional homeostasis.^84–86^ We show that *Tet1* dosage itself acts as a rheostat for circuit reproducibility and binge-eating susceptibility: partial *Tet1* loss converts a normally invariant VTA^DA^ circuit into a probabilistic one. This places *Tet1* within a broader class of epigenetic robustness factors, including *Trim28, Dnmt3a, SetDB1*, and *Nnat*, that regulate phenotypic heterogeneity through transcriptional control.^87–89^ Notably, many of these factors often act across developmental and adult timescales, ensuring stability while preserving plasticity.^64^

*Tet1*-dependent regulation shapes developmental outcomes within the mPFC–VTA axis,^16,90^ indicating that divergent reward setpoints can arise probabilistically from subtle differences in connectivity strength or input balance. Such partially penetrant outcomes are characteristic of multiple neurodevelopmental processes independent of genotype, including callosal agenesis^91,92^ and stochastic defasciculation of the fornix.^93^ Consistent with this interpretation, combined *Tet1/Tet2* deficiency produces variably penetrant neurodevelopmental phenotypes,^94^ as does *Tet1* homozygous deletion.^95^ These observations support a general model in which circuit-level reproducibility and variability coexist within the same developmental program.

A key finding of this study is that the same *Tet1*-dependent machinery that stabilizes circuit assembly during development remains available to modulate circuit output in adulthood. Remodeling of 5hmC has been implicated in reward learning, extinction, and drug conditioning^36,66,96^ suggesting that neuronal activation can re-engage epigenetic pathways originally deployed for developmental canalization. This repurposing of developmental programs for adult plasticity is consistent with the “neural rejuvenation” hypothesis of addiction^97^ and provides a mechanistic framework linking *Tet1* activity to both stable susceptibility and experience-dependent re-calibration. EGR1-directed TET1 activity offers a molecular example of this principle.

Environmental context is likely to influence the balance between epigenetic stability and plasticity.^98^ TET enzymes require vitamin C as a cofactor and operate within folate-dependent one-carbon metabolism, which supplies methyl donors for DNA methylation. Both pathways are essential for neural development and vary across individuals and populations. Vitamin C enhances TET1-mediated hydroxymethylation and dopaminergic differentiation,^99^ whereas folate availability constrains methyl-donor flux.^100^ Variability in these inputs during periods of high embryonic TET1 expression could therefore modulate epigenetic fidelity, increasing variance in VTA connectivity and food reward setpoints without altering genotype. In humans, epigenetic bistability at the TET1 promoter (cg23602092) may similarly provide a substrate for phenotypic diversification, buffering stochastic developmental variation while permitting divergence.^35,75,101^ Although the human analyses rely on peripheral methylation measures, the strong blood– brain correspondence at the TET1 locus, the behavioral associations, and its genetic anchoring support the relevance of these signals to brain epigenetic state. Human analyses were conducted in female cohorts with eating disorders, and future work will be needed to assess generalizability across sexes and diagnostic categories.

Together, these results identify *Tet1*-mediated hydroxymethylation as an epigenetic mechanism that couples developmental robustness with adaptive flexibility. By preserving circuit reproducibility while permitting probabilistic divergence, *Tet1* provides a framework for understanding how behavioral individuality (and susceptibility to binge eating) can emerge from the same developmental logic.

## Methods

### Mice

All mice were bred within the animal facility at the Van Andel Institute (Grand Rapids, MI, USA) and the Max-Rubner Laboratory at the German Institute for Human Nutrition, Potsdam-Rehbrücke (DIfE). Mice were group housed in individually ventilated cages, with *ad libitum* access to chow (mouse breeder diet 5021, #00064, LabDiet, Gray Summit, MO) and water, under a 12-hour on/off light cycle, at a room temperature (RT) of 22 ± 2 °C, with 50–70 % humidity. Whenever indicated, mice had access to high-fat diet (HFD #D12492 with 60 % of calories from fat, Ssniff or Research Diets, New Brunswick, NJ). All experiments were approved by the Animal Ethics Committee of the Van Andel Institute (AUP 22-07-026, PIL-23-04-010, PIL-23-11-028) and the Landesamt für Arbeitschutz, Verbraucherschutz, und Gesundheit (Land Brandenburg, Germany), under applications 2347-20-2021, and conducted in compliance with the ARRIVE guidelines and the EU directive 2010/63/EU. Over the course of this investigation our group became aware of a CNV at the Fgfbp3-Ide locus, present globally in the C57Bl6/J mouse strain to which our transgenic lines are crossed. ddPCR analysis of this CNV did not show any detectable correlation with the heterogeneity phenotypes reported here.

DAT-ires-Cre mice were either directly obtained from Jackson Laboratories (B6.SJL-Slc6a3^tm1.1(cre)Bkmn/J^; JAX 006660) or for fiber photometry experiments provided by the research group of Prof. Nils-Göran Larsson, MPI for Biology of Ageing, Cologne (maintained on a C57BL6/N background). Genotyping of DATCre mice was carried out via PCR using the following primer sequences:

VAI cohorts:

oIMR6625: TGGCTGTTGGTGTAAAGTGG

oIMR6626: GGACAGGGACATGGTTGACT

oIMR8292: CCAAAAGACGGCAATATGGT

WT: 264 bp, Tg: 152 bp.

DIfE cohorts:

DatF3: CATGGAATTTCAGGTGCTTGG

CreR2: CGCGAACATCTTCAGGTTCT

DATR1: ATGAGGGTGGAGTTGGTCAG

WT: 311 bp, Tg: 474 bp.

The PCR cycling conditions were as follows: step 1: 95 °C for 5 min; step 2: 95 °C for 30 s; step 3: annealing 58 °C for 30 s; step 4: 72 °C for 30 s (steps 2–4 repeated for 35 cycles); step 5: 72 °C for 5 min. The resulting PCR products were separated on a 1.5 % agarose gel.

CAG-Sun1/sfGFP mice were obtained from Jackson Laboratories (B6;129-*Gt(ROSA)26Sor*^*tm5(CAG-Sun1/sfGFP)Nat*^/J; JAX (021039) and genotyped using the following primers:

oIMR0872: AAGTTCATCTGCACCACCG

oIMR1416: TCCTTGAAGAAGATGGTGCG

oIMR7338: CTAGGCCACAGAATTGAAAGATCT

oIMR7339: GTAGGTGGAAATTCTAGCATCATCC

WT: 324 bp; Tg: 173 bp.

*Tet1*^fl/fl^ mice^102^ were kindly provided by the research group of Dr. Yong-hui Jiang and genotyped using following primers: 2ndLoxP F1: TGTTGAGAAAAACGGCACTG neoGT F1: TCGACTAGAGCTTGCGGAAC 2ndLoxP R1: GATAGACCACGTGCCTGGAT

WT: 217 bp; Flox: 304 bp.

*Tet1*^+/–^ mice, originally named Tet1^tm1Koh^ and Tet1^tm1.1Koh^ after Cre-mediated excision of the LacZ cassette were obtained from the research group of Dr. Kian Peng Koh.^51^ Mice were crossed according to a heterozygous x wildtype breeding scheme with balanced directionality, i.e., equally distributing transgene origin to either the maternal or paternal allele, respectively. Both male and female mice were metabolically and behaviorally phenotyped. Mice were genotyped using the following primers:

LacZ-tm1-Gtype-6R: CGGATTGACCGTAATGGGATAG

Tet1tm1-Gtype-7F: TTGGCAACACCTCCAGATT

Tet1tm1-Gtype-10R: GCTTTGATGTCTTCGTCTTCATC

WT: 190 bp; Tg: 510 bp.

Tamoxifen-inducible DAT.CreER^T2^ mice (TgSlc6a3/creERT2; MGI: 3835856) were obtained from the research groups of Prof. Cristina Garcia-Caceres and of Prof. Wolfgang Wurst and genotyped using the following primers:

Cre_for: TCTGATGAAGTCAGGAAGAAC

Cre_rev: GAGATGTCCTTCACTCTGATTC

42: CTAGGCCACAGAATTGAAAGATCT

43: GTAGGTGGAAATTCTAGCATCATCC

WT: 324 bp; Tg: 500 bp.

Tamoxifen (Sigma, T-5648) got dissolved in sterile sunflower oil and ethanol (9:1) to a final concentration of 20 mg/mL. At 6 weeks of age, mice were injected with 2 mg of tamoxifen (100ţL; i.p.) once daily for three consecutive days.

Doxycycline-hyclate (D9891, Sigma) got dissolved in drinking water at 2mg/ml and supplemented with 2.5 % sucrose to mask taste. Doxycycline-containing water bottles were provided 2 h before eBED and the first 2 h-limited HFD access time window.

### Surgical procedure

For fiber photometry experiments, analgesia was provided starting two days prior (0.5 mg/mL tramadol in drinking water). Mice were given either buprenorphine (0.1 mg/kg) or meloxicam (5 mg/kg) subcutaneously 20–30 min before surgery. Anesthesia was induced with ketamine + xylazine (100 mg/kg + 7 mg/kg; i.p.) or with 5 % isoflurane in an induction chamber, then maintained at 1–2.5 % via nose cone. After shaving and disinfecting the surgical site, lidocaine gel (2 %) was applied, followed by a rostro-caudal skin incision to expose the skull. After surgery, mice were returned to a heated recovery cage. For the following three days, meloxicam (5 mg/kg; s.c.) was provided bi-daily for pain management. Behavioral testing was performed after 3 weeks of recovery.

### Stereotaxic coordinates

Skull holes were drilled at VTA coordinates (AP −3 mm, ML ± 0.5 mm) and, in some mice, also at mPFC^PL^ coordinates (AP −1.85 mm, ML ± 0.25 mm); for fiber photometry, an additional hole was made for an 1 mm anchor screw close to the frontonasal suture. Using a Hamilton syringe, 300 nL of AAV was injected at 150 nL/min into the VTA (AP −3 mm, ML ± 0.5mm, DV −4.7 mm) or mPFC^PL^ (AP −1.85 mm, ML ± 0.25 mm, DV −1.9 mm).

### Fiber photometry implantation

300 nL of AAV_2/9_-hSyn1-FLEX-GCaMP6s (3.94E13 GC/mL) was injected into the right VTA at 150 nL/min. After scoring the skull with a scalpel for improved adhesion of dental cement, a 1 mm anchor screw was inserted into the rostral skull hole for additional structural support. A 400ţm optical fiber (Doric: MFC_400/430-0.48_-5mm_SM3(P)_FLT) was placed above the right VTA (AP −3 mm, ML +0.5 mm, DV −4.1 mm). Dental cement secured the cannula and screw.

### Fiber photometry recordings

Two separate cohorts of male and female mice (2–4 months old) underwent 7 days of fiber photometry recordings. Of each cohort, mice were randomized to either eBED or cHFD paradigms, respectively. On days 1–2, mice received chow (V1534-300), and on days 2–5 they received 60 % high-fat diet (HFD: Ssniff D12492). In the eBED paradigm, mice accessed HFD for 2 h before returning to chow; in the cHFD paradigm, they had uninterrupted HFD access from first exposure onward (during recordings and home-cage housing).

Two weeks after surgery, mice were habituated for 5 days by being placed in new cages and connected to patch cables for 10 min. During diet intervention experiments, mice were placed in a clean cage with some home-cage bedding, attached to the photometry cable, and acclimated for 10 min. Recordings began with a 10-min baseline (no food), followed by introduction of a glass bowl with pre-weighed food pellets. To prevent excessive photobleaching, recording of VTA^DA^ activity was limited to a total of 60 min. In the eBED paradigm, mice had additional 60 min of HFD access and consumption was calculated from the 2 h difference in food weight. Data were acquired using a LUX RZ10X processor and Synapse software (TDT) with Doric Lenses components. LEDs at 405 nm and 465 nm excited isosbestic and GCaMP6s channels, modulated at 330 and 210 Hz. LED currents were tuned to yield 150–200 mV signals, offset by 5–10 mA, and demodulated with a 6 Hz low-pass filter. Simultaneous behavioral videos were recorded at 10 FPS.

### Fiber photometry analysis

Behavioral Observation Research Interactive Software (BORIS)^3^ was used to manually timestamp the onset of eating events in all behavioural video recordings. The Guided Photometry Analysis in Python protocol (GuPPy)^2^ was used to analyse time-locked behavioral events in the GCaMP6s signal: In brief, any artefacts in the raw signal were removed and the isosbestic control channel (405 nm) was fitted to the signal channel (465 nm) using a least squares polynomial fit of degree 1. Δ*F/F* was then computed by subtracting the fitted control channel *F*_0_ from the signal channel *F* divided by the fitted control channel *F*_0_:

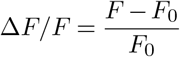

where

*F* = Signal (GCaMP6s 465 nm)

*F*_0_ = Fitted control (Isosbestic 405 nm) .

A standard z-score was calculated from the Δ*F/F* signal:

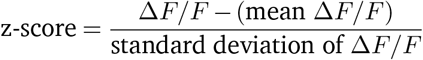

Peri-Stimulus/event Time Histograms (PSTH) of z-score data were generated from −30 to 90 seconds, with a baseline correction between −30 to −10 seconds. Area-under-the-curve (AUC) and Peak z-score values were extracted and used for group-level analysis.

### Chemogenetics

For chemogenetic inhibition, 300 nL of AAV_2/8_-hSyn-DIO-hM4D(Gi)-mCherry (DREADD, Addgene, #44362, 2E13 GC/mL) or AAV_2/8_-hSyn-DIO-mCherry (control, Addgene, #50549, 1.5E13 GC/mL) was injected into the VTA or mPFC^PL^ (with the VTA injected with AAV_rg_-hSyn1-Cre; Addgene, #105553, 2E11 GC/mL)) at 150 nL/min. Three weeks after viral injection, mice were acclimated to metabolic chambers for two days with *ad libitum* access to chow and water. On the following five days, mice received CNO (1 mg/kg; i.p., Tocris) 20 min before 2 h-limited HFD access beginning at dark onset (eBED protocol). All metabolic and behavioral parameters, including binge-style food intake, were recorded throughout the experiment.

### Retrograde input mapping

To perform brain-wide mapping of VTA inputs, AAV_rg_-hSyn1-EGFP (Addgene, #105553, 7E12 GC/mL) was bilaterally injected into the VTA of 6–8-week-old *Tet1*^+/–^ mice and littermate controls. At 3 months of age, mice were exposed to continuous HFD (plus chow) for 4 weeks, followed by 2 days of HFD withdrawal with chow only and the 5-day eBED protocol (2 h-limited HFD access at dark onset). At the end of eBED, mice were transcardially perfused.

### Transcardial perfusion and cryosectioning

At sacrifice, adult mice were either euthanized by CO_2_ asphyxiation or deeply anesthetized (pentobarbital 400 mg/kg; i.p.) and transcardially perfused with phosphate-buffered saline (PBS) followed by 4 % paraformaldehyde (PFA) in borate buffer (pH 8.5). Whole brains were harvested and post-fixed in 4 % PFA for 24 h (or 72 h after fiber photometry) at 4 °C. Brains were then dehydrated in 30 % sucrose in PBS overnight at 4 °C before further processing. Frozen brains were cryosectioned into 40ţm-thick coronal sections using a sliding microtome (PFM Medical Slide 4004 M) or a clinical cryostat (Leica CM1950). Serial sections were either processed immediately or stored in glycerol/ethylene glycol/sucrose containing cryoprotectant at −20 °C, until further use. For retrograde input mapping, every 5^th^ coronal section was selected and subjected to immunostaining (mouse-anti-TH) to label catecholaminergic fibers.

Mouse embryo heads were collected at E15.5 and immersion fixed in 4 % PFA at 4 °C for 48 h and sectioned into 10ţm-thick sagittal sections using a cryostat (Leica CM1950) and directly mounted on microscopic slides.

### Brain Registration

AxioScan .czi images were imported into a QuPath project. The data were organized such that each brain had its own QuPath file. The Aligning Big Brain Atlases (ABBA) plugin through Fiji was used to register each brain to the mouse brain atlas. ABBA can communicate with QuPath; this bridge was used to import all images for one brain, align them, and then export the registrations into the QuPath project. A combination of interactive transformations, Spline transformation, and manual editing of registrations were used. Slices that were missing more than one half of the cortex or severely damaged were excluded from the analysis. The ABBA plugin for QuPath includes a script for importing the brain regions as annotation objects.

### GFP and TH signal area analysis

The whole brain annotation outline (Root object) generated from the ABBA plugin was used to direct automatic threshold calculations. Each Root object was inspected and edited as needed to exclude surrounding background for imperfectly aligned or damaged tissues. This was done to ensure that autofluorescence or other background noise was minimally influential in calculating the threshold value for measuring the GFP or TH signals. Holes within the tissue were not excluded. The Otsu algorithm was used to measure the GFP signal, and the Triangle algorithm was used for the TH signal. The threshold value calculated for the whole brain slice was then used for measuring the area of each brain region covered by either the GFP or TH signal. Custom groovy scripts were used for running these calculations and measurements in batch in QuPath. Area covered by GFP and TH signals in each brain region, calculated threshold values, and area of the brain regions were exported as tsv files for further statistical analysis. Circos plots of input mapping for selected brain regions were plotted using R package `circlize` (V0.4.16).^103^

### Metabolic chambers

Behavioral and metabolic phenotyping was carried out using Promethion metabolic cages (SableSystems, USA) allowing for the measurements of general locomotion and voluntary wheel running activity, as well as indirect calorimetry to calculate energy expenditure and respiratory exchange ratios (RER). Importantly, this setup also allows for the precise assessment of water, chow and HFD intake (featuring automated access doors). Mice were acclimated to the cages for a minimum of 48 h before starting the eBED paradigm. 2 h-limited access was always timed to the beginning of the dark phase. Data was exported and processed using MacroInterpreter (Sable Systems) including the calculation of eating behavior microstructure.

### X-gal staining

X-gal staining was performed as previously described with slight modifications.^104^ In brief, free-floating, 40ţm-thick coronal brain sections (adult) were washed in staining buffer (2 mM MgCl_2_, 0.01 % sodium deoxycholate, and 0.02 % NP-40 in H_2_O) for 10 min at RT. Sections were then incubated with 1 mg/mL X-gal in staining buffer supplemented with 5 mM Potassium Ferricyanide and 5 mM Ferrocyanide at 37 °C overnight.

Embryo head sections (10ţm-thick; sagittal) were mounted on a glass slide first. Staining of slides was carried out as described above but with entire slides being incubated inside a humidified staining chamber at 37 °C overnight.

### Immunofluorescent staining (mouse)

Brain sections were first washed with PBS, pH 7.4, and pre-incubated in 100 mM L-glycine in PBS for 15 min. For 5hmC immunostaining, sections were treated with 1 M HCl for 30 min at room temperature followed by 10 mM sodium citrate buffer, pH 6 + Tween-20 (0.05 % v/v) for 10 min at 95 °C. Then, sections were incubated overnight at 4 °C with primary antibodies (mouse-anti-TH 1:300 + either rabbit-anti-5hmC, 1:300; or rabbit-anti-TET1, 1:300; or rabbit-anti-EGR1, 1:300; or rabbit-anti-cFOS, 1:500) in SUMI (0.25 % porcine gelatine and 0.5 % Triton X-100 in PBS, pH 7.4). Next, sections were serially washed in PBS, pH 7.4, and incubated with respective secondary antibodies (goat-anti-mouse AF488; goat-anti-rabbit AF555; both at 1:1000) diluted in SUMI for 2 h. After another two serial washes in PBS, sections were incubated in DAPI (2ţg/mL in PBS, pH 7.4). Brain slices were mounted and cover slipped (VectaShield HardSet antifade).

Images were acquired as z-stacks using a confocal microscope (Nikon A1plus-RSi Laser Scanning Confocal Microscope or a Leica LSM 880 Laser Scanning Microscope with AiryScan) with an air-immersed 10x and 20x objectives, or oil-immersed 40x and 63x objectives. ImageJ/FIJI, QuPath and IMARIS were used for image processing.

### Immunohistochemistry (human)

Human midbrain sections containing the VTA were cut into 5ţm sections using a Leica RM2235 microtome. Slides were air dried overnight at room temperature followed by a 1 h incubation at 60 °C in Antigen Retrieval and Deparaffinization Buffer using the Dako PT Link in Dako Flex High pH buffer for 20 min at 97 °C. Sections were blocked for 1 h with 0.1 M Tris/2 % FBS (Tris/FBS) prior to overnight incubation at 4 °C in primary antibodies mouse-anti-TH (MABN1188; 1:300) and rabbit-anti-5hmC (Active Motif 39791; 1:500) diluted in Dako Flex Blocking Buffer. After serial washes, sections were incubated in secondary antibodies (goat-anti-mouse AF488 and goat-anti-rabbit AF647; both at 1:500) for 2 h at room temperature followed by DAPI (1:5000) for 10 min.

### Tissue collection

At sacrifice, adult mice were euthanized by CO_2_ asphyxiation and brains were rapidly removed and placed in an ice-cold mouse brain matrix (1 mm; Stoelting Co). Per brain, a coronal slice (Bregma −1.9 to −2.3 mm) was sectioned using razor blades. The VTA was collected using a 1.5 mm micropuncher (Braintree Scientific, USA), immediately flash-frozen on dry-ice, and stored at −80 °C until further processing.

### VTA^DA^ nuclei isolation and Fluorescence-assisted Nuclei Sorting (FANSorting)

CAG-Sun1/sfGFP mice were crossed with DAT-*ires*-Cre mice to generate heterozygous mice. Frozen whole midbrains were individually processed to obtain single nuclei following a previously described protocol.^105^ Frozen midbrains were transferred to a Dounce homogenizer containing 0.7 mL of freshly prepared ice-cold nuclei isolation buffer (0.25 M sucrose, 25 mM KCl, 5 mM MgCl_2_, 20 mM Tris pH 8.0, 0.4 % IGEPAL 630, 1 mM DTT, 0.15 mM spermine, 0.5 mM spermidine, 1x phosphatase & protease inhibitors, and 0.4 units RNasin Plus RNase Inhibitor. Homogenization was achieved by carefully douncing 10 strokes with the loose pestle, incubating on ice for 5 min and further douncing 15 more strokes with the tight pestle. The homogenate was filtered through a 20ţm cell strainer, centrifuged at 1000 xg for 10 min at 4 °C, the nuclei pellet resuspended in 450ţL of staining buffer (PBS, 0.15 mM spermine, 0.5 mM spermidine, 0.4 units RNasin Plus RNase Inhibitor, 0.4 % IGEPAL-630, 0.5 % BSA) and incubated for 15 min on ice. Nuclei pellets were resuspended in 0.5 mL of fresh staining buffer supplemented with DAPI 0.2ţg/ţL. Samples were sorted on a FACSymphony S6 SORP (BD Biosciences) using a 100ţm nozzle with phosphate buffered saline as sheath fluid (IsoFlow Sheath Fluid, Beckman Coulter). Sheath pressure was maintained at 20 PSI and the drop frequency was set at 30 kHz. The 60 mW 349 nm laser was used for DAPI excitation with a 450/50 nm band pass detection filter, and the 200 mW 488 nm laser was used for GFP excitation with a 515/20 nm band pass detection filter threshold was set at 5000 on FSC. Single nuclei were identified on a plot of DAPI-W vs. DAPI-A. Following the nuclei parent gate, GFP+ nuclei were identified on a plot of FSC-A vs. GFP-A and the gate was set and GFP+ nuclei were sorted using a 3+ way sort mask of 0-32-0 (Yield, Purity, and Phase) into 1.5 mL Eppendorf tubes containing (1xDNA/RNA shield; Zymo Research, USA) at RT.

### DNA/RNA isolation

DNA and RNA were isolated from tissues using a commercially available kit (Quick-DNA/RNA Microprep Plus Kit, Zymo Research, USA) and eluted in 16ţL elution buffer (10 mM Tris in H_2_O).

### Reverse transcription and qPCR analysis

Identical amounts of RNA were reverse-transcribed to cDNA using iScript Clear gDNA Synthesis Kit (Bio-Rad, USA) and gene expression was analyzed using TaqMan probes (ThermoFisher Scientific, USA) using 384-well plates (Bio-Rad, USA). Expression changes in *Tet1* (Mm01169087_m1), *Tet2* (Mm01320358_m1), and *Tet3* (Mm01191007_g1) were calculated using the 2^-ΔCt^ method normalized by *Hprt* (Mm03024075) as housekeeping gene.

### Enzymatic Methyl (EM)-seq

Libraries were prepared by the Van Andel Genomics Core. DNA controls: pUC19, unmethylated lambda DNA, and 5hmC T4 (0.008 %, 0.16 %, and 0.12 %) were added to each high molecular weight genomic DNA sample and were sheared to approximately 350 bp average size. Libraries were prepared using the NEBNext Enzymatic Methyl-seq Kit (New England Biolabs, MA) or the NEBNext 5hmc Detection Kit (New England Biolabs, MA) with an input of 2.5 ng of sheared DNA into each prep. Libraries were made according to their respective protocol; briefly, for 5hmc libraries, the NEBNext Post Ligation Supplement was used, and for both preps, the denaturation method used was Formamide, and 11 cycles of PCR were used during amplification. Both Protocols use NEBNext Multiplex Oligos Unique Dual Index Primers Pairs (New England Biolabs, MA). Quality and quantity of the finished libraries were assessed using a combination of Agilent DNA High Sensitivity chip (Agilent Technologies, Inc.), QuantiFluor® dsDNA System (Promega Corp., Madison, WI, USA), and Kapa Illumina Library Quantification qPCR assays (Kapa Biosystems). Individually indexed libraries were pooled, and 150 bp, paired-end sequencing was performed on an Illumina NovaSeq6000 sequencer (Illumina Inc., San Diego, CA, USA) to return a minimum raw coverage depth of 30 × per library. Base calling was done by Illumina RTA3 and output of NCS was demultiplexed and converted to FastQ format with Illumina Bcl2fastq v1.9.0.

### Preprocessing of whole-genome EM-seq data

Raw reads from EM-seq libraries (detecting both 5mC and 5hmC) and 5hmC-only libraries were processed independently using an identical workflow. Briefly, raw FASTQ files were trimmed with Trim Galore (v0.6.10), and the trimmed reads were aligned to the mm10 reference genome, concatenated with FASTA sequences of three spike-in controls, using BISCUIT (v1.4.0).^106^ SAMtools (v1.19)^107^ and dupsifter (v1.2.0)^108^ were used to mark duplicates and to sort and index the resulting BAM files. Subsequently, the `pileup, vcf2bed`, and `mergecg` modules in BISCUIT were used to compute cytosine retention, extract coverage and beta values, and generate methylation calls. The spike-in controls consisted of unmethylated lambda phage DNA, T4 phage DNA containing only 5hmC, and pUC19 plasmid DNA containing only 5mC. Quality control was performed using the QC script included in the BISCUIT package. We evaluated methylation levels and coverage of all three spike-in sequences to confirm that T4 5hmC was detected at comparable levels in both the 5hmC and EM-seq datasets, that lambda phage DNA showed methylation levels near zero, and that pUC19 exhibited full methylation only in the EM-seq libraries.

### Subtraction and differential methylation analysis

Coverage and methylation calls from each sample were used to generate subtraction profiles representing 5mC-specific methylation levels using the `adjustMethylC` function from the R package `methylKit` (v1.30.0).^109^ The subtraction dataset, together with the EM-seq and 5hmC datasets, was then used for differential methylation analysis with the R package `DSS` (v2.25.9).^110^ DSS performs differential methylation detection using a dispersion-shrinkage model followed by Wald statistical testing for each CpG site and genomic region. For differential methylation analysis, CpG matrices were first filtered to retain only sites with coverage greater than 5. Differentially methylated loci (DMLs) were identified using the `DMLtest` and `callDML` functions in DSS with smoothing enabled. Differentially methylated regions (DMRs) were subsequently identified using the `callDMR` function with parameters delta = 0.05 and p.threshold = 0.01.

Genome annotations for the mm10 reference were retrieved using the `getAnnot` helper function from the package `DMRseq` (v1.24.1),^111^ which accesses annotation resources provided by `annotatr` (v1.30.0).^112^ Visualization of methylation data and DMRs was performed using the `plotDMRs2` function from the R package `DMRichR` (v1.7.8).^111,113,114^

### Analyses in the ESTRA and IMAGEN datasets

#### Cohorts

##### Participants with eating disorders

A total of 74 individuals with eating disorders were included from the ESTRA study, comprising 34 participants with bulimia nervosa (BN; mean age = 22.4 ± 2.13 years) and 40 participants with anorexia nervosa (AN; mean age = 22.0 ± 2.16 years).^68^ All participants met DSM-5 diagnostic criteria for BN or AN, as assessed using the Eating Disorder Diagnostic Scale (EDD).^115^ Participants were recruited through the Eating Disorders Unit at the South London and Maudsley National Health Service Foundation Trust or via targeted social-media advertisements.

##### Healthy controls

Seventy-two healthy control (HC) participants (mean age = 20.1 ± 2.27 years) were included, comprising 19 individuals recruited within ESTRA and 53 age- and sex-matched individuals from the IMAGEN cohort.^69,116^ All participants were female, aged 18–25 years, of European ancestry, and recruited in London. Healthy controls were younger than participants with AN or BN (both *p* < 0.05); therefore, age was included as a covariate in all downstream analyses.

Binge-eating frequency was assessed in all participants using the item “number of binge eating days per week”. Written informed consent was obtained from all participants prior to study participation.

#### DNA methylation profiling and preprocessing

Whole-blood samples were from ESTRA and IMAGEN participants and processed jointly. Genome-wide DNA methylation (DNAm) was quantified using the Illumina Infinium HumanMethylationEPIC v2 BeadChip, interrogating over 900 000 CpG sites, following the manufacturer’s standard protocols.

Raw intensity data (IDAT) files were imported into R using the `minfi` package, generating RGChannelSet objects containing red and green channel intensities. Beta values, representing the proportion of methylated signal at each CpG site, were extracted using the `getBeta` function.

Sample-level quality control (QC) was conducted on the combined dataset prior to normalization. Initial QC assessment used the `minfi qcReport` function to evaluate signal intensity distributions and Illumina control probes. Beta values were then derived following background correction and internal control normalization using `preprocessIllumina`, restricted to autosomal probes. Multidimensional scaling was performed on a random subset of 10 000 CpG sites, and principal component analysis (PCA) was conducted using singular value decomposition of centred beta values. Samples exceeding ± 3 standard deviations from the median on any of the first four methylation principal components were classified as outliers and excluded.

Sex was predicted from sex-chromosome methylation patterns using the `minfi getSex` function (cutoff = −2), and samples with discrepancies between predicted and recorded sex were removed. Methylation control samples, when present, were identified via sample-sheet annotations or predefined identifiers and excluded. All flagged samples were removed, and the raw methylation data were reloaded to generate a cleaned RGChannelSet for downstream processing.

Following sample-level QC, quantile normalization was applied using the `minfi preprocessQuantile` function with stratified normalization, fixing outliers and automatically removing samples with low signal intensity that could not be corrected (log_2_ median methylated or unmethylated intensity below the default threshold of 10.5). Normalised beta values were extracted and restricted to autosomal probes for all downstream analyses.

Probe-level QC was then applied. Probes mapping to the X or Y chromosomes were excluded. CpG sites were removed if they showed poor technical performance (detection *P* values > 0.01 in > 20 % of samples); for remaining probes, failed measurements were set to missing. Probes affected by common genetic variation were excluded, including those with single-nucleotide polymorphisms at the CpG site, single-base extension site, or within the probe body (minor allele frequency > 5 %). Probes known to be cross-reactive or to hybridise to multiple genomic locations were also removed based on published annotations. Finally, probes showing invariant methylation across samples (defined as beta values ≤ 0.2 or ≥ 0.8 in all samples) were excluded.

To account for cellular heterogeneity, proportions of major blood cell types were estimated using a reference-based approach implemented in `minfi` with the `FlowSorted.Blood.EPIC` reference panel. Estimated proportions of CD8^+^ T cells, CD4^+^ T cells, natural killer cells, B cells, monocytes, and neutrophils were included as covariates in downstream analyses. Residual technical variation was further controlled by including the first four DNA methylation principal components derived from the normalized beta-value matrix.

All preprocessing and QC scripts are publicly available at:

https://github.com/XinyangYu918/DNA-methylation-preprocessing.

#### Genotype and methylation quantitative trait loci (meQTL) mapping

Participants were genotyped using the Illumina Human610-Quad BeadChip, Human660-Quad BeadChip or Illumina Global Screening Array v3.0. Full details of genotype QC have been described previously^117^ and are also available at

https://github.com/XinyangYu918/

EatingBehaviours-BrainMaturation-

Psychopathology-Genetics.

Briefly, stringent QC was applied prior to imputation, and participants identified as ancestry outliers relative to the European superpopulation were excluded. Genotype data passing QC were imputed using the European ancestry reference panel from the 1000 Genome Project (phase 3, release v5).

Cis-meQTL mapping was performed using the `MatrixEQTL` package.^118^ Linear regression models were fitted to test associations between genotype and DNAm levels, adjusting for age, batch effects, the first four multidimensional scaling (MDS) components derived from genotype data, the first four DNA methylation principal components, and estimated blood celltype proportions. Significant meQTLs were validated using published cis-meQTL summary statistics from an independent adolescent cohort.^72^

#### fMRI monetary incentive delay (MID) task

Participants completed a modified MID task during functional MRI to assess neural responses to reward anticipation and reward outcome. The task comprised 66 trials (10-s each). At trial onset, one of three cue shapes was presented for 250 ms, indicating potential reward magnitude (0, 2, or 10 points) and target location (left or right). Following a variable anticipation period (4000–4500 ms), participants were instructed to respond as quickly as possible to a target stimulus (white square) via button press. Target duration (100–300 ms) was dynamically adjusted to achieve an individual success rate of approximately 66 %. Feedback indicating reward outcome (number of points were won, if any) was displayed post-response for 1450 ms.

For the present analyses, contrasts of interest were anticipation of high reward versus no reward, and feedback of high reward versus no reward. Only trials with successful target hits were included.

Region-of-interest (ROI) analyses focused on the dorsomedial prefrontal cortex (dmPFC). Blood-oxygen-level–dependent (BOLD) responses during reward anticipation and feedback were examined in relation to binge-eating frequency, DNA methylation at cg23602092 (located within the *TET1* promoter region), and rs2664444, the identified meQTL. All models were adjusted for age, diagnostic group, and scanner site. Methylation-based analyses additionally included the first four DNA methylation principal components and estimated blood cell-type proportions, while genetic analyses included batch and the first four genotype-derived MDS components.

### Analyses in other human datasets

#### Blood–brain DNA methylation correspondence

Blood–brain DNA methylation correspondence was evaluated using a publicly available database derived from matched whole-blood and post-mortem brain samples [Epigenetics 2015; 10(11): 1024-1032; https://epigenetics.essex.ac.uk/bloodbrain/]. The dataset includes 144 samples from 122 individuals, with DNA obtained from whole blood and four brain regions (prefrontal cortex, entorhinal cortex, superior temporal gyrus, and cerebellum). Correlations between blood and brain methylation levels at CpG sites within the human *TET1* promoter were assessed to estimate cross-tissue concordance.

#### Brain expression quantitative trait loci (eQTLs)

The associations between rs2664444 and brain gene expression were obtained from the GTEx Portal (accessed 03/11/2025).

## Author Contributions

T.G. and J.A.P. conceptualized all studies and designed all experiments. T.G. (stereotaxic surgeries, metabolic and behavioral phenotyping, immunohistochemistry, imaging and analyses, qPCR), T.G. and R.C. (fiber photometry), X.Y. and H.B. (human eating disorder cohort methylome and fMRI), L.F., S.A., and Z.F. (bioinformatic analyses), K.G. (image analyses), M.DA. (HPLC analysis of brain monoamine metabolites), R.V., M.H., J.G., H.D., T.C., and L.DC. (experimental support) conducted the experiments, collected, and analyzed the data. T.G. and J.A.P. wrote the manuscript in discussion with R.N.L., K.T., and S.D., who revised the article critically for important intellectual content. All authors have read and approved the final version of the manuscript.

## Acknowledgments

The authors thank Anika Desphande for her assistance. The authors thank the Van Andel Institute Genomics Core (RRID:SCR_022913), especially Katelyn Becker, Mark Wegener, Tracy Avequin, and Dr. Marie Adams, for providing facilities and assistance with EM-seq, and the Van Andel Institute Bioinformatics and Biostatistics Core (RRID:SCR_024762), especially Dr. Zhen Fu, for her assistance with EM-seq analyses. We thank the Van Andel Institute Pathology and Biorepository Core (RRID:SCR_022912), especially Lisa Turner, for her assistance with tissue processing and immunostaining of human post-mortem samples. This research was supported in part by the Van Andel Institute’s Brain Bank (MiND; RRID:SCR_-026035); Optical Imaging Core (RRID:SCR_021968), especially Dr. Kristin Gallik, Dr. Lorna Cohen, Dr. Jian Tai, and Dr. Corinne Esquibel; Flow Cytometry Core (RRID:SCR_022685), especially Kohl Sprader, Maddie Nichols, and Dr. Rachel Sheridan; and Transgenic and Vivarium (RRID:SCR_022914 and RRID:SCR_-023211), especially Megan Tompkins, Reta Burdette, William Weaver, Audra Guikema, Adam Rapp, and Scott Bechaz. This work was supported in part by funding to T.G. from the Alexander-von-Humboldt Foundation (Feodor-Lynen Postdoc Fellowship) and the German Center for Diabetes Research DZD (Post-doc Travel Grant); from VAI through internal philanthropy to J.A.P., National Institutes of Health awards R01HG012444 to J.A.P. and J.H.N., R01DK132216 to J.A.P; the MRC and Medical Research Foundation (ESTRA—Neurobiological underpinning of eating disorders: integrative biopsychosocial longitudinal analyses in adolescents: Grant No. MR/R00465X/ to S.D.; ESTRA—Establishing causal relationships between biopsychosocial predictors and correlates of eating disorders and their mediation by neural pathways: Grant No. MR/S020306/1 to S.D.), and was co-funded by UK Research and Innovation under the U.K. government’s Horizon Europe funding guarantee (Grant Nos. 10041392 and 10038599) as part of the Horizon Europe HORIZON-HLTH-2021-STAYHLTH-01 (Grant Agreement No. 101057429: environMENTAL to S.D.). This paper represents independent research, partly funded by the National Institute for Health and Care Research (NIHR) Maudsley Biomedical Research Centre (BRC). The views expressed are those of the author(s) and not necessarily those of the NIHR or the Department of Health and Social Care. Funding support for R.N.L. and R.C. on the project was provided by the Leibniz Association through the Leibniz Competition Best Minds Grant “BAByMIND” (J99/2020) and the Deutsche Forschungsgemeinschaft (DFG, German Research Foundation) under Germany’s Excellence Strategy – EXC-2049 – 390688087 (NeuroCure) to R.N.L.). This work was partially funded by the European Union within the scope or the European Research Council ERC-CoG Trusted no. 101044445, awarded to T.D.M. T.D.M. further received funding from the German Research Foundation (DFG TRR296, TRR152, SFB1123 and GRK 2816/1) and the German Center for Diabetes Research (DZD e.V.)

## Consortia

### ESTRA Consortium

Lauren Robinson, Marina Bobou, Zuo Zhang, Gareth J. Barker, Gunter Schumann, Ulrike Schmidt, Sylvane Desrivières.

### IMAGEN Consortium

Tobias Banaschewski, Gareth J. Barker, Arun L. W. Bokde, Christian Büchel, Herta Flor, Antoine Grigis, Hugh Garavan, Penny Gowland, Andreas Heinz, Jean-Luc Martinot, Marie-Laure Paillère Martinot, Frauke Nees, Dimitri Papadopoulos Orfanos, Luise Poustka, Michael N. Smolka, Henrik Walter, Robert Whelan, Sylvane Desrivières, Gunter Schumann.

### PERMUTE Consortium

J. Andrew Pospisilik, Ilaria Panzeri, Luca Fagnocchi, Stefanos Apostle, Emily Wolfrum, Zachary Madaj, Jillian Richards, Holly Dykstra, Tim Gruber, Mitch McDonald, Andrea Parham, Brooke Armistead, Timothy J. Triche Jr., Zachary DeBruine, Mao Ding, Ember Tokarski, Eve Gardner, Joseph Nadeau, Christine Lary, Carmen Khoo, Ildiko Polyak, Qingchu Jin.

## Data availability

The data that support the findings of this study are available from the corresponding author upon reasonable request. For the human cohort data (ESTRA and IMAGEN), please contact Sylvane Desrivières.

## Supplementary Material

**Supplementary Figure 1.**
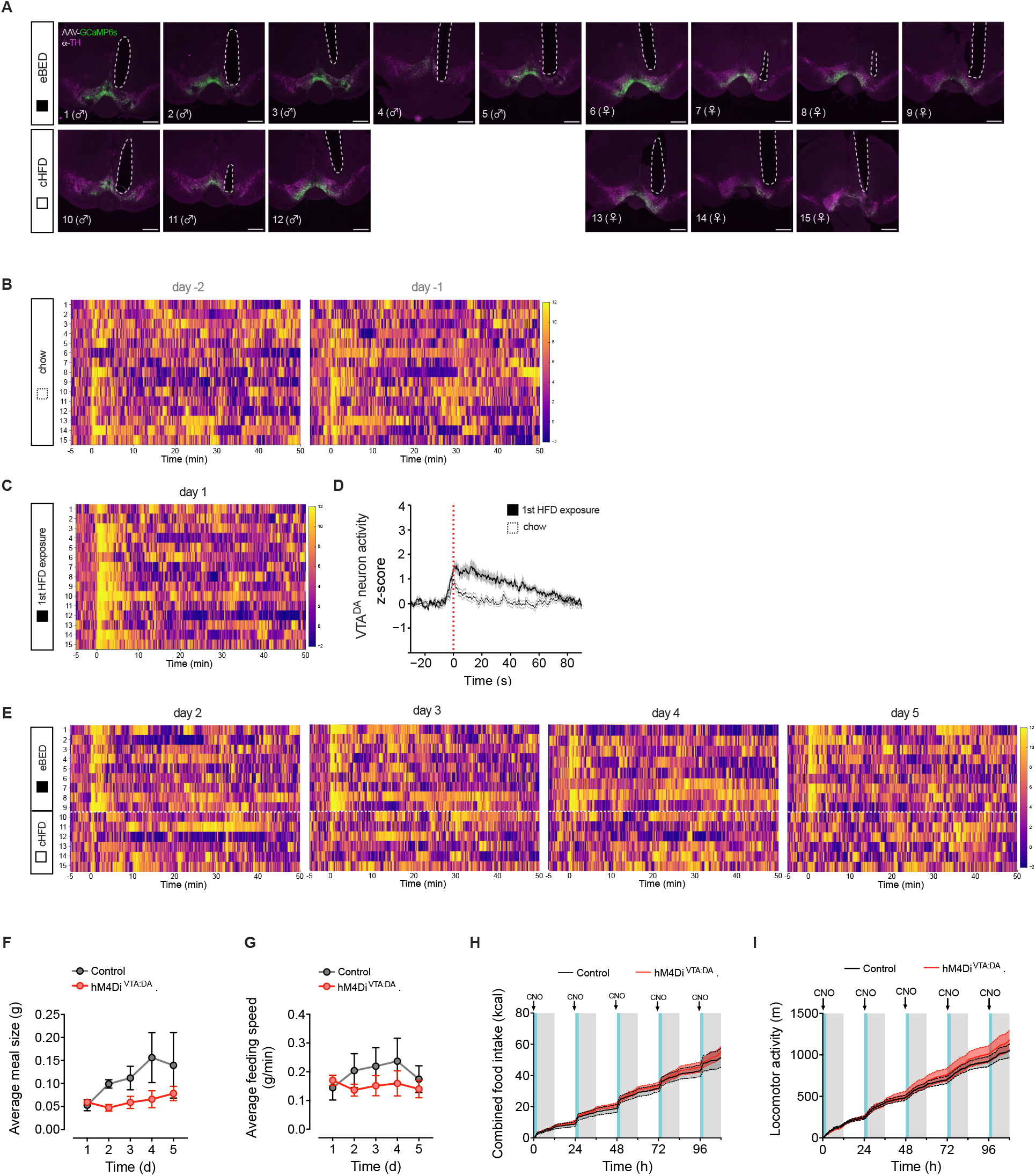
Related to Figure 1. Escalatory VTA^DA^ activity is triggered by eBED and required for binge eating in mice. **(A)** Targeting validation of fiber photometry experiment including all mice; GCaMP6s expression colocalizing with VTA^DA^ neurons (TH^+^) and optic fiber tract, respectively. **(B)** Heatmap representation of fiber photometry *z*-scores of individual mice over the course of the entire recording sessions on day −2 and day -1 (chow diet; food given at *T* = 0 min). *n* = 6–8 mice. **(C)** Heatmap representation of fiber photometry *z*-scores of individual mice over the course of the entire recording sessions on day 1 (first HFD exposure; food given at *T* = 0 min). *n* = 6–8 mice. **(D)** Fiber photometry VTA^DA^ recordings of mean *z*-score of the first HFD exposure versus the two baseline recordings (chow diet) at eating onset (0 min). Data are presented as mean ± SEM. *n* = 6–8 mice. **(E)** Heatmap representation of fiber photometry *z*-scores of individual mice over the course of the entire recording sessions on day 2–5 (eBED versus cHFD; food given at *T* = 0 min). *n* = 6–8 mice. **(F)** Average meal size per day of hM4Di^VTA:DA^ mice relative to control mice. Data are presented as mean ± SEM. *n* = 4–7 mice. **(G)** Average eating speed per day of hM4Di^VTA:DA^ mice relative to control mice. Data are presented as mean ± SEM. *n* = 4–7 mice. **(H)** Cumulative combined food intake (HFD + chow) of hM4Di^VTA:DA^ mice relative to control mice. Time window with limited HFD access (shaded) and CNO injection (1 mg/kg BW; i.p.; arrow) are indicated. Data are presented as mean ± SEM. *n* = 4–7 mice. **(I)** Cumulative locomotor activity of hM4Di^VTA:DA^ mice relative to control mice. Time window with limited HFD access (shaded) and CNO injection (1 mg/kg BW; i.p.; arrow) are indicated. Data are presented as mean ± SEM. *n* = 4–7 mice.

**Supplementary Figure 2.**
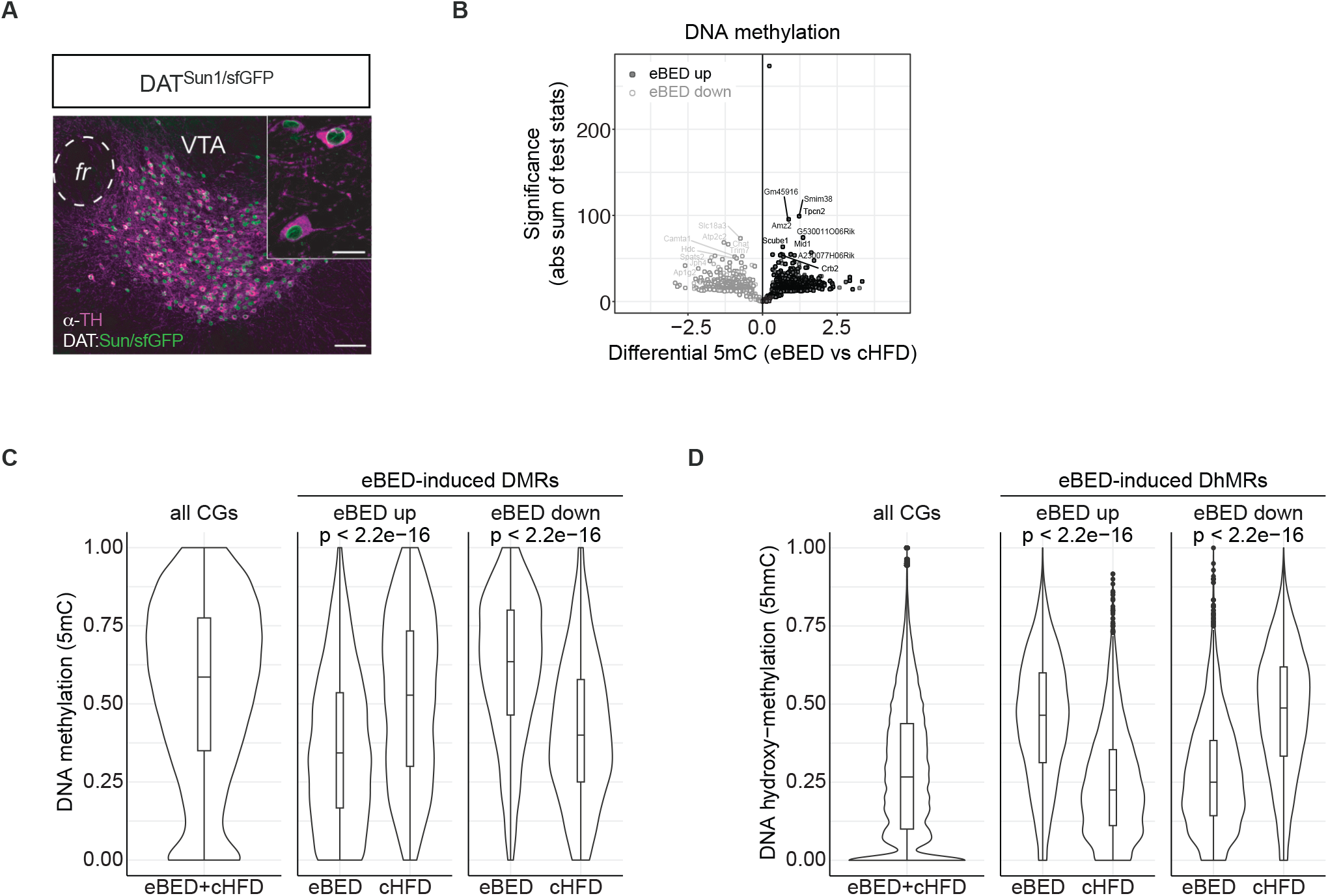
Related to Figure 2. eBED triggers rapid and profound rewiring of the VTA^DA^ neuron epigenome. Confocal micrograph of DAT^Sun1/sfGFP^ reporter mice with GFP-tagged VTA^DA^ nuclei; scale bars: 50 ţm, 20 ţm (insert). Volcano plot of DMRs comparing eBED versus cHFD. **(C)** Violin plots showing the levels of 5mC on all eBED+cHFD CGs (reference) and eBED-induced DMRs, showing decreased (left) or increased (right) amount of the DNA mark with respect to cHFD. *P* -values from Wilcoxon test. **(D)** Violin plots showing the levels of 5hmC on all eBED+cHFD CGs (reference) and eBED-induced DhMRs, showing increased (left) or decreased (right) amount of the DNA mark with respect to cHFD. *P* -values from Wilcoxon test. In all panels, DMRs’ effect size cut-off = 0.05; *p*-value cut-off = 0.01 from Wald statistical testing. *N* = 12 (2 samples each condition — eBED, cHFD, chow — and each mark -5mC, 5hmC).

**Supplementary Figure 3.**
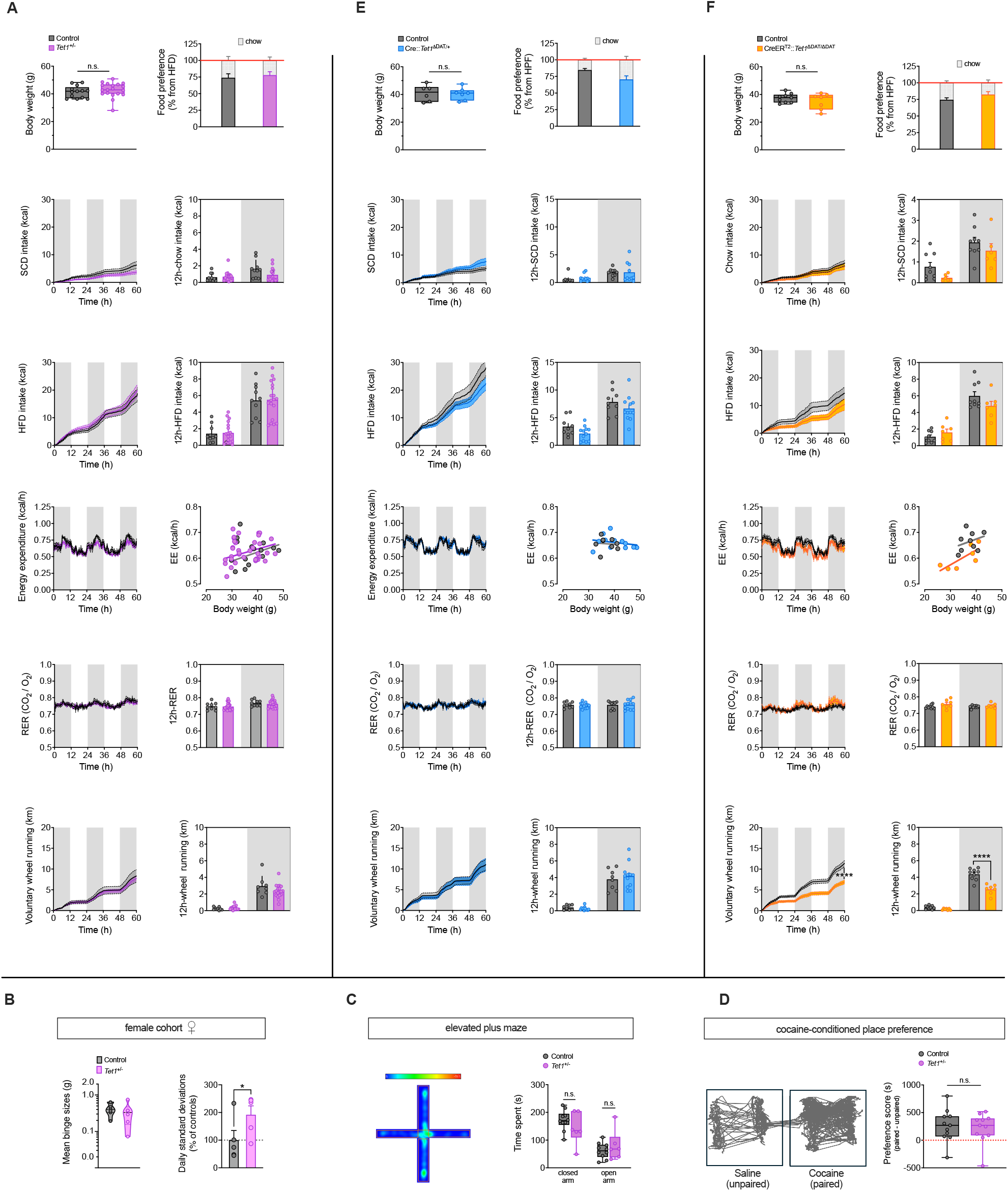
Related to Figure 3. Heterozygous loss of *Tet1* triggers heterogeneity in binge-eating vulnerability. **(A)** In-depth metabolic phenotyping of male *Tet1*^+*/*−^ mice. *n* = 10–22 mice. **(B)** Mean binge sizes of *Tet1*^+*/*−^ female mice and female littermate controls (left panel), plotted on a Log10 scale (shaded area: normal binge response). Data are presented as violin plots with median and quartiles. *n* = 6–7 mice. Standard deviation of daily binge sizes after Log10 transformation relative to respective controls (right panel). Data are presented as mean ± SEM of STDEV from Day 1–5. ∗*P <* 0.05 (one-tailed Student’s *t*-test). **(C)** Elevated plus maze (EPM) performed on male *Tet1*^+*/*−^ mice and controls with representative movement tracking (left panel) and corresponding quantification (right panel). Data are presented as median ± minimum and maximum. *n* = 6–10 mice. **(D)** Cocaine-conditioned place preference performed on male *Tet1*^+*/*−^ mice with representative movement tracking (left panel) and corresponding quantification (right panel). Data are presented as median ± minimum and maximum. *n* = 11 mice. **(E)** In-depth metabolic phenotyping of male Cre::*Tet1*^ΔDAT/+^. *n* = 9–12 mice. **(F)** In-depth metabolic phenotyping of male iCreER^T2^::*Tet1*^ΔDAT/ΔDAT^. *n* = 7–9 mice.

**Supplementary Figure 4.**
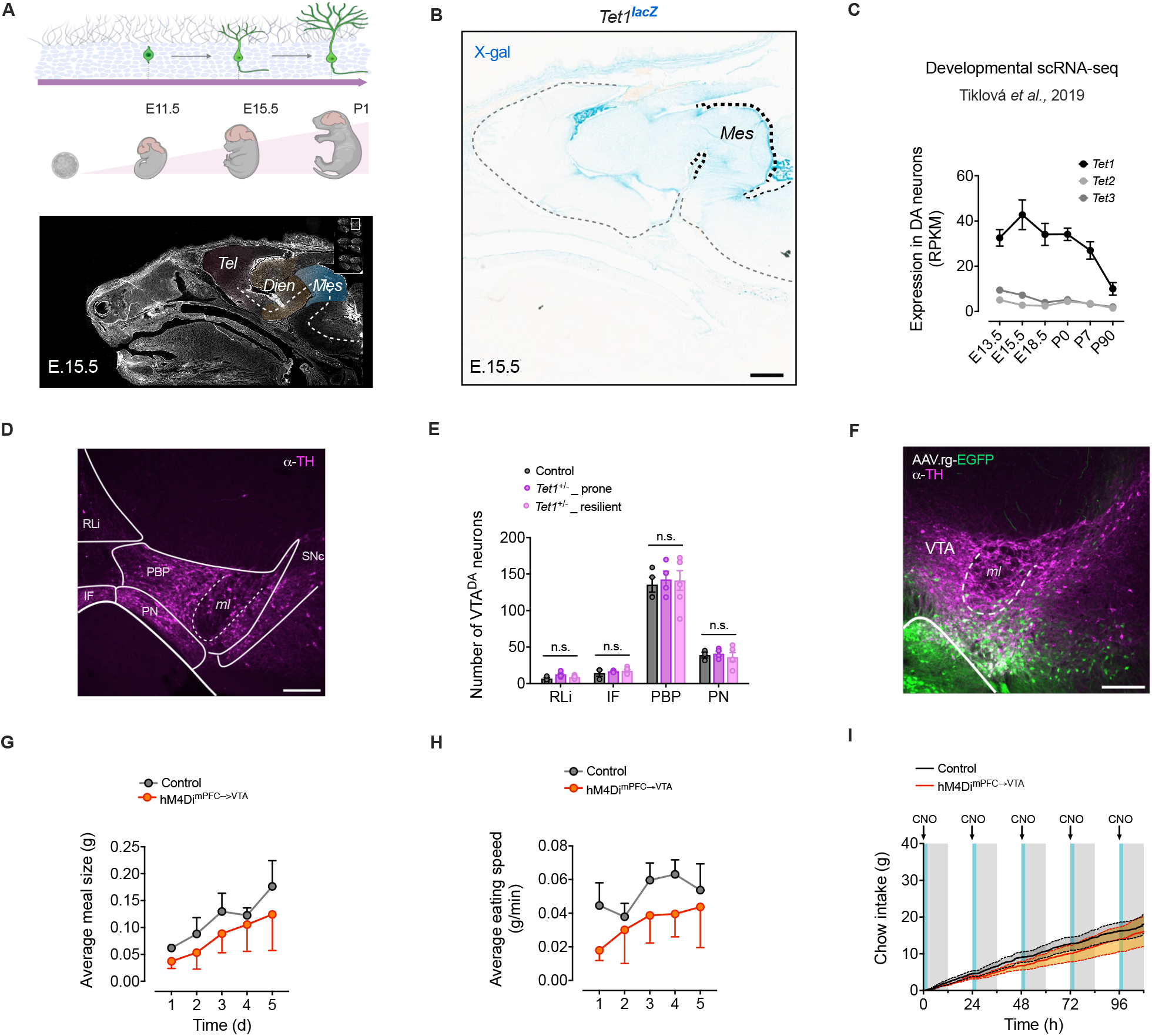
Related to Figure 4. Reduced mPFC^PL^→ VTA inputs confer binge-eating resilience in *Tet1*^+*/*−^ mice. **(A)** Sagittal overview of embryo head (E15.5) with pseudocolored brain regions (Telencephalon: red; Diencephalon: yellow; Mesencephalon: blue). *Created in BioRender.com*. **(B)** *Tet1*^*LacZ*^ reporter mouse embryo (E15.5) after X-gal staining (Mesencephalon highlighted). Scale bar: 500 ţm. **(C)** Reanalysis of developmental scRNA-seq^58^ showing expression levels of *Tet1, −2* and *−3* at different embryonic stages of development. **(D)** Micrograph showing VTA^DA^ neurons (magenta; TH^+^) across different VTA subregions of an adult male mouse. **(E)** Quantification of VTA^DA^ neuron counts per VTA subregion. **(F)** Micrograph of injection site showing VTA^DA^ neurons (magenta; TH) and EGFP^+^ interneurons and fibers. **(G)** Average meal size per day of hM4Di^mPFC-VTA^ mice relative to control mice. Data are presented as mean ± SEM. *n* = 4–7 mice. **(H)** Average eating speed per day of hM4Di^mPFC-VTA^ mice relative to control mice. Data are presented as mean ± SEM. *n* = 4–7 mice. **(I)** Cumulative chow intake of hM4Di^mPFC-VTA^ mice relative to control mice. Time window with limited HFD access (shaded) and CNO injection (1 mg/kg BW; i.p.; arrow) are indicated. Data are presented as mean ± SEM. *n* = 4–7 mice.

**Supplementary Figure 5.**
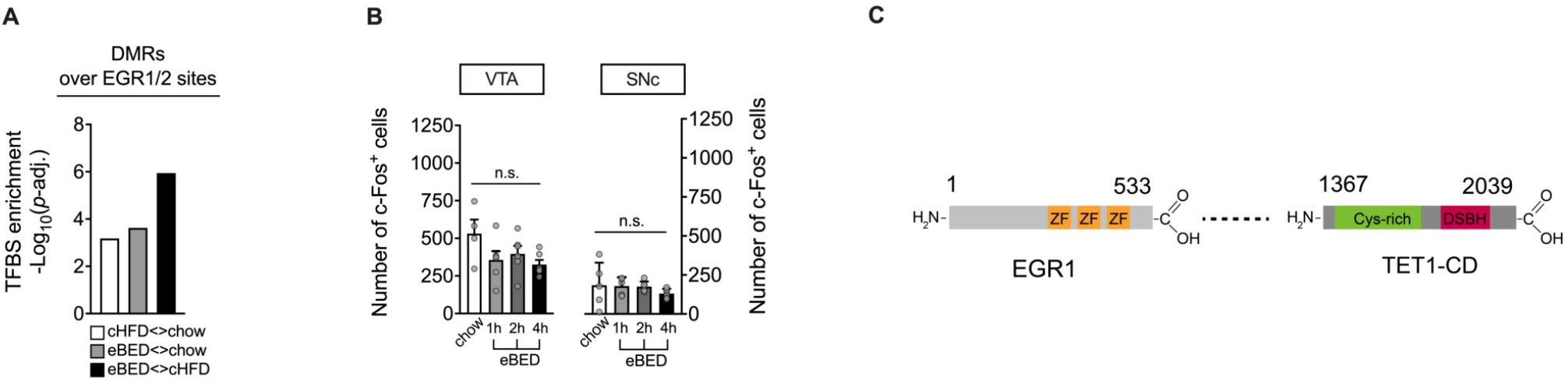
Related to Figure 5. EGR1-guided TET1 reactivation in VTA^DA^ neurons restores binge-eating susceptibility. **(A)** Motif analyses on eBED-induced DMRs showing significant enrichment of indicated transcription factors, including EGR1. Adjusted *p*-values from one-tailed Fisher’s exact test, followed by Bonferroni correction. **(B)** Quantification of c-Fos immunoreactivity punctae in the VTA and SNc at 1 h, 2 h or 4 h after the last of five binge-eating episodes versus chow-fed control mice. Data are presented as mean ± SEM. n.s. = not significant. *n* = 4–7 mice (one-way ANOVA). **(C)** Schematic illustration of EGR1-TET1.CD fusion protein. ZF: zinc finger domain, Cys-rich: cystine-rich domain, DSBH: double-stranded *β*-helix domain.

**Supplementary Figure 6.**
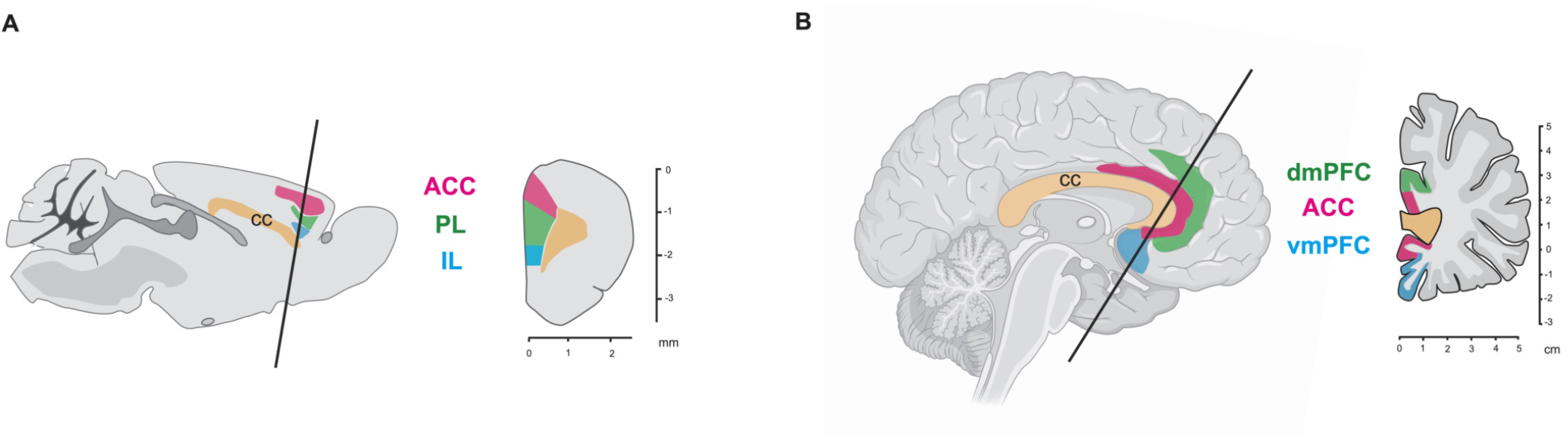
Related to Figure 6. Human *TET1* methylation status mediates binge eating and dmPFC brain activation during reward processing. **(A)** Illustration of a mouse brain in sagittal (left) and coronal (right) orientation highlighting medial prefrontal cortex (mPFC) subregions. ACC: anterior cingulate (Cg1) mPFC, PL: prelimbic mPFC, IL: infralimbic mPFC. *Created in BioRender.com*. **(B)** Illustration of a human brain in sagittal (left) and coronal (right) orientation highlighting medial prefrontal cortex (mPFC) subregions. dmPFC: dorsomedial PFC, ACC: anterior cingulate mPFC, vmPFC: ventromedial PFC. Homologous regions are color-coded (green: PL ↔ dmPFC; red: ACC; blue: IL ↔ vmPFC).

